# Taxonomy, nomenclature, and identification of the giant hummingbirds (*Patagona spp*.) (Aves: Trochilidae)

**DOI:** 10.1101/2024.07.03.601580

**Authors:** Jessie L. Williamson, Chauncey R. Gadek, Bryce W. Robinson, Emil Bautista, Selina M. Bauernfeind, Matthew J. Baumann, Ethan F. Gyllenhaal, Peter P. Marra, Natalia Ricote, Nadia D. Singh, Thomas Valqui, Christopher C. Witt

**Affiliations:** Cornell Lab of Ornithology, Cornell University; Ithaca, 14850, USA; Department of Ecology and Evolutionary Biology, Cornell University; Ithaca, 14850, USA; Museum of Southwestern Biology, University of New Mexico; Albuquerque, 87131, USA; Department of Biology, University of New Mexico; Albuquerque, 87131, USA; Environmental Stewardship, Los Alamos National Laboratory; Los Alamos, 87545, USA; Centro de Ornitología y Biodiversidad (CORBIDI); Lima, 15064, Peru; The Earth Commons Institute; Department of Biology; McCourt School of Public Policy, Georgetown University; Washington, D.C., 20057, USA; Facultad de Artes Liberales, Departamento de Ciencias, Universidad Adolfo Ibáñez; Santiago, 7941169, Chile; Department of Biology, Institute of Ecology and Evolution, University of Oregon, Eugene, 97403, USA; Facultad de Ciencias Forestales, Universidad Nacional Agraria La Molina; Lima, 15024, Perú

**Keywords:** Andes, bird, biodiversity, cryptic species, genus revision, identification key, intraspecific variation, nomenclature, morphological comparison, taxonomic revision

## Abstract

Giant hummingbirds (*Patagona* spp.) are extraordinarily large hummingbirds whose taxonomy has been muddled for two centuries. *Patagona* systematics were recently redefined in a study of migration, physiology, and genomics, revealing two species: the Southern Giant Hummingbird and Northern Giant Hummingbird. Here, we re-evaluate taxonomy and nomenclature of the genus in light of its newly-clarified biology and species limits, analyzing data from 608 specimens and wild-caught individuals spanning 1864–2023. The forms *gigas* and *peruviana* were both described based on multiple syntypes. No adequate syntypes for *P. gigas* are extant, so we designate a neotype for this taxon. We then critically consider the identity and usage of *gigas* and *peruviana*, respectively, and examine identification challenges that have fostered taxonomic uncertainty. We endorse the names *Patagona gigas* for the Southern Giant Hummingbird and *P. peruviana* for the Northern Giant Hummingbird. The genetic identity of the *peruviana* lectotype remains untested, but its plumage appears to match the northern species. We found that ∼33% of *Patagona gigas* specimens in major museum collections are misidentified as *peruviana*; we provide this list to correct the historical record. To facilitate identification and future study of these two cryptic species, we provide comprehensive information on plumage, measurements, and seasonal ranges.

## INTRODUCTION

Giant hummingbirds (*Patagona spp.*) comprise the most distinctive clade in the family Trochilidae because of their extraordinary size. Within the genus *Patagona*, however, morphological and plumage variation are subtle and it is difficult to disentangle geographic variation from age, wear, molt, sex and/or individual variation. Seasonal overlap among cryptic but distinct geographic breeding populations has exacerbated the challenge of characterizing the differences among genetic lineages and appropriately defining taxonomy.

A recent study made it possible to accurately adjudicate taxonomy within *Patagona* for the first time by presenting new information about genetics, migration, and phenotypes (Williamson et al. 2024; Fig. 1). We now know that *Patagona* is a clade with two biological species that diverged ∼2–3 Mya and that differ in breeding latitude and migratory behavior. The two lineages are largely allopatric during the breeding season. During the austral winter, the southern lineage, which breeds from central Chile through northwestern Argentina and likely throughout Bolivia, migrates to tropical latitudes of the central Peruvian Andes where it overlaps extensively with the northern, high-elevation resident lineage. The difficulty of distinguishing the two cryptic species based on plumage and/or measurements has, until now, made it difficult to characterize their seasonal range overlap. However, the two lineages are reproductively isolated and experience no gene flow, making their status as biological species clear (Williamson et al. 2024). Each of the two species, the Northern Giant Hummingbird and Southern Giant Hummingbird, contains shallow geographic population structure. The high-elevation resident Northern Giant Hummingbird averages larger in bill length, wing chord, tail length, and tarsus length, although there is substantial overlap in morphological characteristics with the migratory Southern Giant Hummingbird––only ∼65–85% of individuals can be identified by measurements when comparing sexes separately. The two species have near-identical body mass distributions. Similarly, subtle plumage differences between the northern and southern species are useful for identification but not universally applicable across the range of individual variation (Fig. 1).

**Figure 1.**
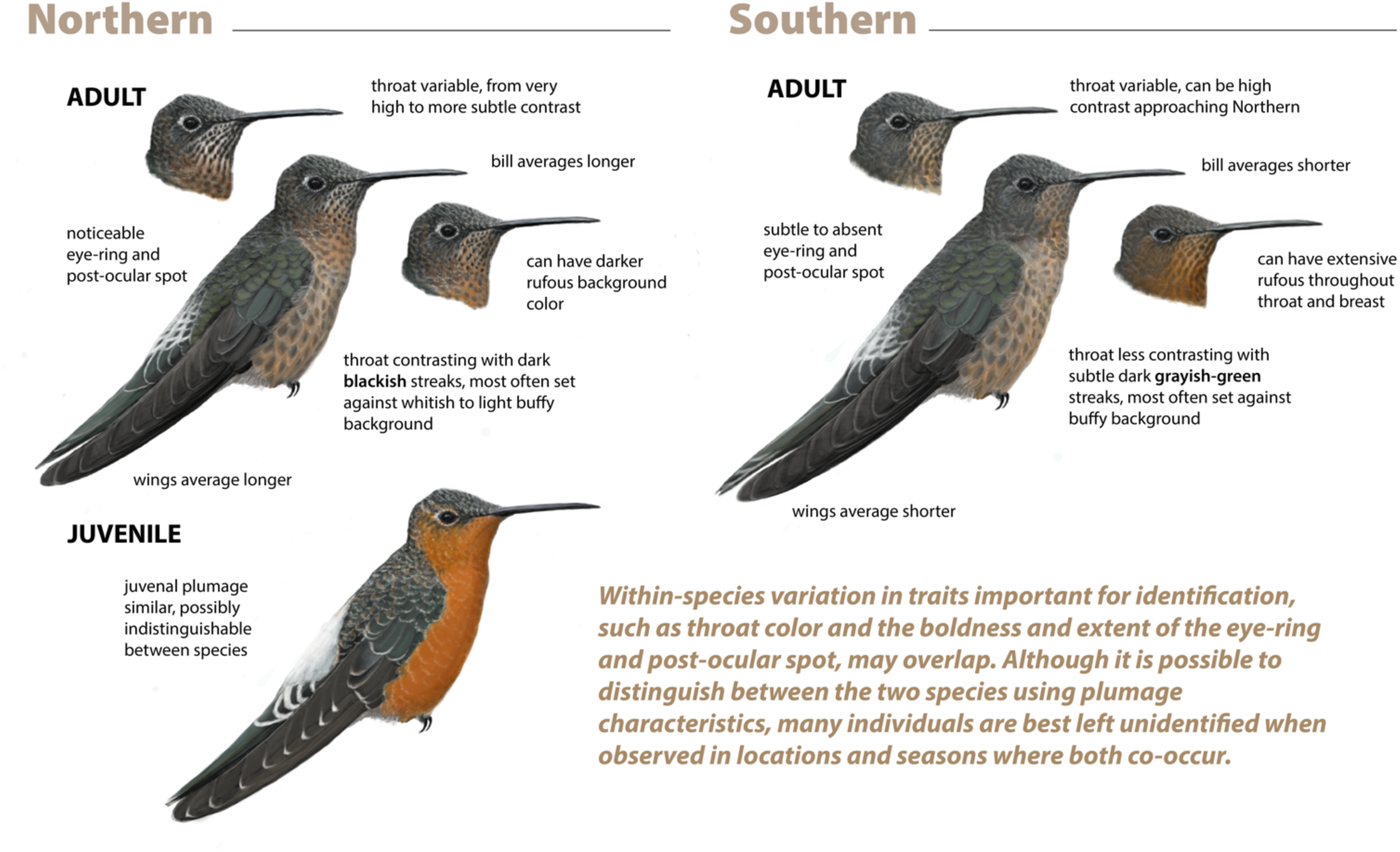
Key for distinguishing Northern and Southern Giant Hummingbirds, highlighting important plumage characteristics described in Williamson et al. (2024), including throat color and pattern and extent of eye-ring and post-ocular spot, as well as morphological characteristics, such as wing length and bill length. Juvenile is represented for comparison (the term “juvenal” is a specific term corresponding to juvenile plumage). Illustrations: Bryce W. Robinson.

With unambiguous species status and newly defined taxon limits, Williamson et al. (2024) proposed a new species name for the Northern Giant Hummingbird, picaflor gigante del norte in Spanish: *chaski* (*Patagona chaski* sp. nov.) and designated a holotype. The name *peruviana* had been introduced by Boucard (1893) in a brief description, and has since been applied at the subspecies level to all giant hummingbirds found from northwestern Argentina and Bolivia northward. Williamson et al. (2024) argued that the name *peruviana* should not continue to be used for two main reasons: (1) Historic and continued misalignment of the name *peruviana* with the northern genetic lineage; and (2) The type series (syntypes) appeared to have been comprised of a mix of the two giant hummingbird species.

However, the taxonomic description by Williamson et al. (2024), overlooked key information regarding the syntypes of *peruviana* that came to the attention of the authors only after publication; an omission that we rectify here. Namely, Williamson et al. (2024) did not know that the likely identities of syntypes used by Boucard (1893) to designate *peruviana* were known. Although Boucard did not provide identifiers for the specimens he referenced in his description, he stated that they were “…collected by Whitely in Peru” (Boucard 1893). At least 16 specimens were collected by Henry Whitely, Jr. in southern Peru from 1867–1873 or later (Supplement; Table S1). Williamson et al. (2024) were not aware of two specimens still at the Muséum National d’Histoire Naturelle in Paris (MNHN-ZO-MO-1989-340 and MNHN-ZO-MO-1989-341) which are considered to be Boucard’s (1893) *peruviana* syntypes and are labeled accordingly (e.g., Zimmer 1930, Hellmayr 1932, Jouanin 1950; Fig. S1a-b; Table S1). These syntypes appear to correspond to the Northern Giant Hummingbird based on plumage features, although the female specimen is somewhat atypical. The identifications of both should be considered tentative until confirmed with DNA sequencing. Shortly after Williamson et al. (2024) was published, Areta et al. (2024) designated MNHN-ZO-MO-1989-340 as a lectotype for *peruviana*. This quick taxonomic action by Areta et al. (2024) sought to affirm the link between the Northern Giant Hummingbird and the name *peruviana*.

A guiding purpose of the International Code of Zoological Nomenclature (the Code) is “to promote stability and universality in the scientific names of animals and to ensure that the name of each taxon is unique and distinct”, thereby ensuring precision and consistency in the use of terms (ICZN 1999). Williamson et al. (2024) stated that they proposed the species name *chaski* because the name *peruviana* had been misapplied to a mixture of two species, both the Northern and Southern Giant Hummingbird, and that the species identities of the specimens listed in the description were dubious. Because the newly-designated *peruviana* lectotype appears to correspond to the Northern Giant Hummingbird, the names *chaski* and *peruviana* should be considered synonyms, with *peruviana* taking precedent because it is senior. Art. 23.1 of the Code, the Principle of Priority, states the general principle that “the valid name of a taxon is the oldest name applied to it…” (ICZN 1999).

Here, we present the taxonomic history and nomenclature of *Patagona* from 1824 to present, elaborating on Williamson et al. (2024) and Areta et al. (2024) and correcting errors in both papers. We review new information about species limits, genetics, migration, and identification that clarifies seasonal ranges and systematics of this genus. We additionally evaluate the first species description by Vieillot (1824) and that of the genus *Patagona* Gray 1840, and examine those of subsequently introduced taxon names, *peruviana* and *boliviana* by Boucard (1893), as well as the specimens that were key to historical treatments and nomenclature. Finally, we designate a neotype to fix the identity of the Southern Giant Hummingbird.

## MATERIALS AND METHODS

### Abbreviations

AMNH, American Museum of Natural History, New York, USA; ANSP, Academy of Natural Sciences of Drexel University, Philadelphia, USA; CM, Carnegie Museum of Natural History, Pittsburgh, USA; CORBIDI, Centro de Ornitología y Biodiversidad, Lima, Peru; CUMV, Cornell University Museum of Vertebrates, Ithaca, USA; FMNH, Field Museum of Natural History, Chicago, USA; LSUMZ, Louisiana State University of Museum of Natural Science, Baton Rouge, USA; Museum of Comparative Zoology at Harvard University, Cambridge, USA; MNHN, Muséum National d’Histoire Naturelle, Paris, France; MSB, Museum of Southwestern Biology, Albuquerque, USA; NHMUK, Natural History Museum, Tring, UK; USNM, Smithsonian National Museum of Natural History, Washington, D.C., USA.

### Taxonomic history of *Patagona gigas*

The species *Trochilus gigas* was described by Vieillot (1824) based on two syntypes from “Brésil” (Brazil) (Vieillot 1824). Vieillot wrote that one specimen “…appears to us to be only a young male; the old one is unknown to us; but we believe we recognize the female in an individual in the possession of Mr. Becœur, naturalist…” (Vieillot 1824). Lesson (1829) renamed the species *Ornismya tristis*, noting that its “homeland” was Chile (Lesson 1829). In 1840, Gray named the genus *Patagona*, listing *P. gigas* (ascribed to Vieillot, 1824) as the sole species (Gray 1840). In spite of other proposed names (e.g., *Hylocharis gigas*, *Ornismya gigantea*, *Trochilus tristis*, *Cynanthus tristis*, and *Hypermetra gigas*; see Elliot 1879, Porter 1897, Hartert 1900), from this point forward, *Patagona gigas* entered into widespread use among ornithologists.

Vieillot’s putative “young male” syntype, illustrated by Paul Louis Oudart in Plate 180 of *La Galerie des Oiseaux* and attributed to M. (Mr.; for “monsieur” in French) Portier of the Ministry of the Navy (Vieillot 1824; Fig. S2) has not been located and its current whereabouts are unknown to us. Vieillot’s putative female syntype is believed to be NHMUK-1933.11.14.36, a taxidermied mount at NHMUK (Warren 1966; Fig. S3). This specimen was obtained by George Loddiges from Jean-Baptiste Bécouer, a pharmacist and collector known for his arsenical soap method of specimen preparation, who died in 1777 (Rookmaaker et al. 2006). It is not known whether NHMUK-1933.11.14.36 was determined to be a female during dissection or whether sex was presumed based on plumage (the latter would be unreliable). This putative female syntype bears two sets of confusing labels that make its identity as a syntype tentative; neither label is original, and both were written at different times and in different hands (date and author unknown): One is a green tag that appears to state “Trochilus gigas Vielliot female / gal. ois. 1.1e 29 p. 296 pl 180 / Loddiges Coll / May be type of female” (note misspelling of “Vieillot”; Fig. S3). The other tag, a white British Museum label, reads on the front, “Trochilus gigas female / Loddiges Colln. (Purchased)” and on the back states, “1933-11-14-36. / may be. See Type catalogue” (again note misspelling of “Vieillot”). The line through the text “Not accepted as Viellot’s type of female but” was done in blue ink at what appears to be a later date (Mark Adams, *pers. comm.*; Fig. S3). Because specimens from the Loddiges collection, purchased by NHMUK in 1933, bore no original labels, it is not possible to verify that they specimens are, indeed, the type specimens of the species they represent; thus, Loddiges specimens are listed by NHMUK as probable holotypes or syntypes (Warren 1966).

Despite the unknown collection dates and provenance (including both specific locality and higher geography) of Vieillot’s two putative syntypes, much later, in 1932, Austrian ornithologist and neotropical bird expert Carl Hellmayr suggested that the (syn)type locality of Brazil described by Vieillot (1824) was an “errore”, stating “…we suggest Valparaíso, Chile” (Hellmayr 1932). On pg. 231 of the *Birds of Chile*, Hellmayr justified his type locality restriction, writing, “Oudart’s plate of *T. gigas*, based on an evidently immature bird from “Brésil” in the collection of ‘M. Portier, attaché au ministère de la marine,’ while none too good, corresponds fairly well to certain bright-colored Chilean specimens, such as No. 61,676, Caldera, and accordingly I propose to restrict Vieillot’s term to the small southern form suggesting Valparaiso [, Chile] as type locality” (Hellmayr 1932). Hellmayr stated that Oudart’s plate illustrated an “immature” bird, but it actually depicts a juvenile (immature birds are subadults which differ in plumage from juveniles and resemble adults in measurements and plumage). The specimen “No. 61,676” referenced by Hellmayr is FMNH-61676, an immature female from Caldera, Atacama Region, Chile, corresponding in plumage to the Southern Giant Hummingbird (we confirmed this by examining the specimen at FMNH). Thus, Hellmayr’s decision to restrict the specific (syn)type locality for *Patagona gigas* to Valparaíso, Chile, was based on his comparison of an immature specimen (FMNH-61676) from Atacama Region, Chile (unconnected to the syntypes), with the Oudart’s illustration of a juvenile putative male from an unknown locality. This restriction of the type locality based on visual evidence was unjustified by any known geographic variable trait.

Further confusing the identity of putative (syn)types, in 1893, Adolph Boucard, a French ornithologist and specimen trader, wrote, “I have in my collection what I consider as the type of Vieillot ‘Ex Coll Riocour’” (Boucard 1893). Boucard references obtaining Vieillot’s type specimen from the Comte de Riocour collection, which he procured most of in 1889 (Renshaw 1905). The specimen he mentions is MNHN-ZO-2023-396 (sex and locality unknown; collection date prior to 1842) at the MNHN, which bears multiple tags, including Comte de Riocour tags with characteristic red outlines and Boucard Collection tags with characteristic purple outlines (Fig. S4). The Comte de Riocour tag reads, “Ex. Coll Riocour / Oiseau mouche geánt / *Trochilus giganteus* / Vieill”. The front side of the Boucard Collection tag reads, “Patagona gigas / Ex. Col. Riocour Type Vieillot ?” Boucard’s tag includes a question mark (“?”) after the words “type Vieillot” (Patrick Bousses, *pers. comm.*), indicating doubt about whether the specimen was, indeed, a Vieillot (syn)type; Boucard seems to have additionally indicated this doubt in his text by writing “…what I consider the type…” (Boucard 1893). However, we could find no evidence that MNHN-ZO-2023-396, which Boucard believed was “Vieillot’s type”, is *the* type or even a syntype of *P. gigas*, nor could we find any indication that this specimen was ever examined by Vieillot. Vieillot described *P. gigas* based on two syntypes: the “young male” (whereabouts unknown) from M. (Mr.) Portier of the Navy and a putative adult female (likely NHMUK-1933.11.14.36), obtained by Jean-Baptiste Bécouer and then by George Loddiges before being purchased by NHMUK (Warren 1966). The specimen MNHN-ZO-2023-396 was obtained by Boucard from the Comte de Riocour collection and resides at the MNHN. There is no evidence that Boucard (or Comte de Riocour before him) ever had access to George Loddiges’ collection, which remained within the Loddiges family after George Loddiges’ death in 1846 (Mark Adams, *pers. comm.*). Consistent with this history, the specimen NHMUK-1933.11.14.36 does not bear original Boucard Collection tags, which would be expected had this specimen been “in my collection”, as Boucard described.

Thus, the entity formerly known as the Giant Hummingbird (*Patagona gigas*) and currently known as the Southern Giant Hummingbird (*Patagona gigas, sensu* Williamson et al. 2024) was described based on a putative type series consisting of two to three presently unconfirmed syntypes with unknown collection dates, localities, and provenance, and whose history is uncertain.

### Taxonomic history of *peruviana* and *boliviana*

In 1893, Boucard proposed the names *Patagona peruviana* and *Patagona boliviana* for specimens in his collection, “…if the two should prove distinct species” (Boucard 1893). Boucard wrote, “…I have also three specimens collected by Whitely in Peru, and in Bolivia by Buckley” (Boucard 1893; Supplement; Fig. S1). In total, Henry Whitely collected at least 16 giant hummingbird specimens (not the 13 mentioned in Areta et al. 2024; Table S1). Boucard’s original text does not list specific identifiers, collection dates, collection localities, or sexes for the specimens in his collection from Chile, Peru, or Bolivia. However, two Whitely specimens at the MNHN (MNHN-ZO-MO-1989-340, a male from Tinta, Peru collected 15 June 1868 and MNHN-ZO-MO-1989-34, a female from Tinta, Peru with no date), were assigned equal status as *peruviana* syntypes, while MNHN-ZO-MO-1989-342 (undetermined sex from Bolivia with no date), collected by Clarence Buckley, was designated as the holotype for *boliviana* (Jouanin 1950; Fig. S1; Table S1). The names *peruviana* and *boliviana*, in addition to the nominate *gigas*, were henceforth recognized and entered into use (e.g., Hartert 1900).

In 1921, Boucard’s (1893) original description was contested by Eugène Simon, a French naturalist and systematist from the MNHN. Simon wrote that *Patagona gigas* is a bird of the high mountains “…without presenting sufficient local variation to be considered as subspecies” (Simon 1921). Simon stated that Boucard (1893) had “…conditionally proposed the name *Patagona peruviana* for the specimens from Peru from Whitely’s trip and *Patagona boliviana* for those brought back from Bolivia by Buckley”. He then added, “…I have ascertained, by examining the types, that the coloration characters [Boucard] gives (in Gen. Humm. Birds. p. 61) are age-related or even completely illusory” (Simon 1921). In one footnote, Simon noted the white borders on the wing feathers and wing coverts and “the absence of a rufous part in front of the white uropygial band” on Boucard’s types, which are characteristics of *Patagona* juveniles and immatures (see Fig. 1). In a second footnote, Simon clarified his description of the phrase “completely illusory”, noting that he was discussing, “…for example, the partly black throat of Peruvian birds” (Simon 1921). Simon did not provide specific identifiers of the supposed types, but it is presumed that he examined the same specimens referenced by Boucard (1893).

In 1930, American ornithologist John Zimmer provided a brief account of *Patagona gigas peruviana* Boucard and *Patagona gigas boliviana* Boucard in *Birds of the Marshall field Peruvian expedition 1922-1923* (Zimmer 1930), recognizing Boucard’s names as subspecific identifiers. Zimmer described examining four recently-collected *Patagona* specimens from Peru, compared against 23 previously-collected Peru specimens and 13 *gigas* specimens from central and northern Chile and northwest Argentina. Zimmer wrote, “The Peruvian birds, with one or two exceptions, are consistently larger than the Chilean skins, with longer wings and longer, heavier bills. They also, with one or two exceptions, are more rufescent on the under parts, sometimes markedly so. Since Dr. Hellmayr plans to discuss the characters of these two races in a forthcoming paper on the birds of Chile, I will leave further details to his able pen” (Zimmer 1930).

Two years later, Hellmayr published *The Birds of Chile* (Hellmayr 1932). In his account for the giant hummingbird, he wrote, “Although the late Eugène Simon (Hist. Nat. Troch., p. 157) questioned the possibility of discriminating any geographic races of the Giant Humming bird, the study of between fifty and sixty properly labeled specimens from the whole range clearly indicates the existence of two forms” (Hellmayr 1932). Hellmayr said Simon’s assessment that *P. peruviana* and *P. boliviana* were *nomina nuda* was a mistake, but Hellmayr agreed with Simon when he noted “…several of the characters claimed by Boucard prove to be individual” (Hellmayr 1932). Hellmayr went on to describe plumage differences among populations from Chile compared to those from Ecuador, Peru, and Bolivia, noting that the extent and intensity of rufous color to the underparts seemed to be “purely individual” and unrelated to sex or age (Hellmayr 1932). Hellmayr concluded, “While the palest examples of the northern form can be closely matched by one or two unusually rufous-bellied birds from central Chile, the general run of the two series is easily told apart” (Hellmayr 1932). His firm conclusion seemed to leave no doubt that there were two forms that were easily distinguishable.

Based on his assessment, Hellmayr offered nomenclatural suggestions: “The larger northern race is entitled to the name *P. gigas peruviana*, tentatively proposed by Boucard for a specimen from Peru in his collection. The type, a male obtained by H. Whitely on June 15, 1868, at Tinta, Dept. Cuzco, agrees with the average of our Peruvian series, while *P. boliviana*, was based on an individual variant with wholly cinnamon-rufous under parts…” (Hellmayr 1932). Hellmayr thus cast doubt on *boliviana*, while buttressing *peruviana*. Hellmayr noted that “…An adult male from Tilcara, Jujuy (Nov. 24, 1905; L. Dinelli), and a couple from Lara, Tucuman (Feb. 12, 1903; G. A. Baer), all in the Tring Museum, are in every particular typical of the large northern form” (Hellmayr 1932). Years later, Hellmayr added that specimen no. 868/48 collected by L. Dinelli on August 10, 1916, from Colalao del Valle, Tucumán, Argentina was “typical” of *peruviana* based on plumage and morphology (Revilliod et al. 1942).

Hellmayr’s 1932 account is largely regarded as the definitive taxonomic treatment of *Patagona* in the literature. Based on his plumage and morphological analysis and nomenclatural suggestions, subsequent authorities recognized *peruviana* as a valid subspecies and ceased recognizing *boliviana*, folding *boliviana* into subsequent treatments of *P. g. peruviana* (Peters 1945, Zimmer 1952)––despite the uncertainty surrounding the distribution of forms.

In 1952, John Zimmer revisited *Patagona* taxonomy in *Studies of Peruvian Birds* (Zimmer 1952). In his account of *P. g. peruviana*, he stated, “This northern form of the species is fairly readily determinable in distinction from the southern *gigas gigas* when fully adult birds are compared, but young examples can be confusing” (Zimmer 1952). Zimmer went on to describe the difficulty of distinguishing juvenile birds by plumage. He noted, “All the adult examples from [the Urubamba Valley of Peru] are clearly *peruviana*. I have seen no Peruvian specimens that indicate the occurrence of *gigas gigas* in Perú, although there is evidence that the subspecies has some migratory tendencies, as noted by Hellmayr…” (Zimmer 1952). Zimmer presented a list of 83 specimens that he examined from numerous localities and of both sexes. He wrote, “I would emend a statement made by Hellmayr (*loc. cit.*) with reference to certain Argentina specimens that he found to be ‘in every particular typical of the large northern form.’ Three females from Tilcara are fairly typical *peruviana* but the others are identifiable only by their measurements, since they are immature and in coloration resemble other young of both forms” (Zimmer 1952). Later, in 1974, Ecuadorian biologist and naturalist Fernando Ortiz-Crespo supported previous scholars, noting “[The race *peruviana*]…differs from the nominate race, occurring in central Chile, adjacent Argentina, and the southern Chilean provinces, in having a larger body size, a stouter, longer bill, and rufescent underparts” (Ortiz-Crespo 1974).

Thus, since ca. 1930, the giant hummingbird has been considered to contain two forms: The nominate *P. g. gigas*, a suspected migrant and supposed smaller-bodied form with a shorter bill and shorter wings, stated to range from approximately central Chile to central Argentina; and *P. g. peruviana*, a supposed larger-bodied form with a longer bill and longer wings, thought to range (and breed) from northwestern Argentina north through Bolivia, northern Chile, Peru and Ecuador. Individuals of *peruviana* were believed to have more rufescent underparts (although previous workers debated about whether the rufous coloration might be related to age, sex, and/or individual variation). The two forms were described as being “easily distinguishable” based on plumage and morphology. It is notable that numerous historical treatments have parallels with the results of modern genomic and tracking studies, despite those historical scholars having no access to DNA sequences. However, the early treatments also introduced errors and greatly oversimplified the phenotypic distinction between the two species, resulting in misidentification, errant inference of ranges, and failure to recognize seasonal range overlap (see below).

### Historical characterization of migration

The migration of Chilean giant hummingbirds has been suspected and discussed since at least 1834, but every aspect of the migratory journey had remained a mystery ever since. Throughout history, naturalists puzzled over where Chilean populations “disappeared” to after breeding (Goodall et al. 1946). Even recently, the *Atlas of the Breeding Birds of Chile 2011-2016* called the winter whereabouts of Chilean-breeding giant hummingbirds an “unsolved mystery” (Medrano et al. 2018). This paucity of migration data has contributed to longstanding confusion about the ranges, taxonomy, and identification of the two giant hummingbird forms.

Darwin was among the earliest to note migration of the Southern Giant Hummingbird: During his voyage on *The Beagle*, he remarked, “At Valparaíso, in the year 1834, I saw several of these birds in the middle of August, and I was informed they had only lately arrived from the parched deserts of the north” (Darwin 1839). German-Chilean botanist Federico Johow wrote that giant hummingbirds are “rather tropical” and were observed in central Chile only during the Chilean summer; he noted that their presence along the coast of Chile coincided with the flowering of *Lobelia* and columnar cacti, proposing that they wintered in the northern provinces of “…Peru, Ecuador, etc.” (Johow 1910, 1921). Hellmayr (1932) stated that *Patagona* is absent from central Chile during June–August, writing, “The birds probably migrate northwards, but a few, at least, cross the Andes on their migration, as is shown by specimens… which clearly pertain to the Chilean race” (Hellmayr 1932). Concurring with this hypothesis, *Birds of the High Andes* posited that Chilean breeders migrate across the Andes to adjacent western Argentina (Fjeldså and Krabbe 1990), while others have suggested that they migrate to northwestern Argentina (Sánchez Osés 2003). More recently, the guide “*Biodiversidad de Chile, patrimonio y desafíos*” asserted without evidence that Chilean giant hummingbirds migrate to Ecuador for the austral fall (Ambiente 2008).

In 1952, Chilean naturalist Rafael Barros offered what is perhaps the most detailed early treatment of *Patagona* migration: Barros described Chilean-breeding giant hummingbirds as “…fleeing from the rainy season and the cold” (Barros 1952). He wrote, “It is not well elucidated if, when it leaves its places of residence…it actually moves to the provinces located to the north of Aconcagua (where my observations begin), or it crosses the high mountain range of the Andes and passes to the Republic of Argentina, or if it goes to winter in some regions of Bolivia or Peru. I know of no observations in this regard, and this is a problem to be solved, of great scientific interest” (Barros 1952). Barros posited the question: “Does it go up…to the heights of the Puna to cross the Andes?”, later responding: “Based on what I have been able to observe for many years… I do not believe that it crosses the mountain range in the central part of the country, during its pilgrimages, either on its outward journey or on its return, because in the mountainous region it first reaches the lower parts, the pre-cordillera, the foothills of the Cordillera and fields beneath them, gaining altitude afterwards” (Barros 1952). This same year, Zimmer cast doubt on the extent of northward travel of southern temperate breeders, stating that he had seen no Peruvian specimens suggestive of *P. gigas gigas* (Zimmer 1952).

In 1974, Fernando Ortiz-Crespo published a detailed treatment of the ecology of giant hummingbirds in Ecuador. He noted that the giant hummingbird population in central Chile “…builds up during the spring and summer”, attributing their movement to northward migration (Ortiz-Crespo 1974). Ortiz-Crespo credited Johow (1910) and Darwin (1839) as his sources for migration information, both of whom had previously characterized Southern Giant Hummingbird migration as unresolved. In this same study, Ortiz-Crespo analyzed morphology between sexes and subspecies (which he assigned based on collection country), stating that his results “…do not seem to confirm the possibility that Ecuador is regularly visited on migration by birds which nest in central Chile, and whose winter quarters thus remain to be discovered” (Ortiz-Crespo 1974). Twelve years later, in a different manuscript, Ortiz-Crespo expressed continued uncertainty about *Patagona* migration, stating “…it is not known for sure where they winter” (Ortiz-Crespo 1986). He described examining 180 specimens, including 156 specimens from outside of Chile, only four of which (ANSP-145417 and FMNH-179394 from Bolivia, ANSP-167558 from Argentina, and FMNH-67518 from Peru) he said had measurements and coloration of the nominate form *gigas* (we examined these same specimens at ANSP and FMNH, and have confirmed that they are Southern Giant Hummingbirds). Ortiz-Crespo concluded, “This meager evidence suggests that southern populations disperse northward after nesting” (Ortiz-Crespo 1986), providing no additional specifics. He further suggested that examining specimens collected during the austral fall and winter from northwest Argentina and Bolivia would be important for understanding distribution and movement (Ortiz-Crespo 1986). Thus, although *Patagona* migration had been discussed in the literature since ∼1834, every aspect remained unknown before Williamson et al. (2024).

### Re-evaluating historical identification criteria for *peruviana*

Hellmayr’s (1932) text is regarded as definitive for *Patagona* taxonomy, particularly with regard to distinguishing *gigas* from *peruviana*. In it, Hellmayr examined morphological characteristics (bill, wing, and tail lengths) from “…between fifty and sixty properly labeled specimens from the whole range…”. Hellmayr concluded that there are “clearly” two forms and that “…the general run of the two series is easily told apart” (Hellmayr 1932). In his text, he presented a table of measurements from 45 specimens from Ecuador (no localities), Peru, Bolivia (one specimen; no locality), Argentina, and Chile. He suggested that *P. g. gigas* was markedly smaller and did not overlap morphologically with *P. g. peruviana*, supporting his conclusion that “the general run of the two series is easily told apart” (Hellmayr 1932). This claim was reinforced by Areta et al. (2024), who stated that Hellmayr “provided a diagnosis of *peruviana* on plumage and morphometry” that was “clearly mirrored” by the data provided in Williamson et al. (2024).

However, Hellmayr’s conclusions about the ease with which *gigas* and *peruviana* could be distinguished contradict results of analyses by Ortiz-Crespo (1974) and Williamson et al. (2024). Among 361 *Patagona* specimens from Ecuador, Peru, Bolivia, Argentina, and Chile, Williamson et al. (2024) found substantial inter-species overlap in the same three morphological characteristics studied by Hellmayr (Williamson et al. 2024). Northern and Southern Giant Hummingbirds differed significantly, on average, in bill length, wing chord, and tail length, respectively; however, average trait differences were extremely subtle, on the scale of millimeters (e.g., ∼1–5 mm)––size differences between the two species were not typically distinguishable by eye, and the ranges of trait values for both species overlapped substantially (Williamson et al. 2024). Similarly, within each giant hummingbird species, sexes differed significantly––but not substantially––in nearly all measurements, despite lack of sexually dimorphic plumage. When analyzing morphological data separately for each sex, linear discriminant analyses using data from three traits (bill, wing, and tail) combined could only distinguish adult female Northern from Southern Giant Hummingbird in 65.4–82.8% of cases and adult males in 79.5–85.7% of cases (Williamson et al. 2024). The mass ranges of the two species are near-identical (Williamson et al. 2024), indicating general body size equivalence.

To understand Hellmayr’s (1932) conclusion that the two forms are easily distinguished, we compared Hellmayr’s original data (*n*=45 adults) with published data from Williamson et al. (2024), measurements from wild-caught Northern and Southern Giant Hummingbirds captured in Peru during July–August 2023, and data from specimens measured at ANSP, CM, FMNH, and USNM during June and August 2024 (*n*=608 individuals total). All individuals were measured by [ANONYMIZED AUTHOR INITIALS]. After removing juveniles and considering only individuals of known sex, locality, and age, our dataset included *n*=448 adults.

We predicted that if Hellmayr’s analysis of *peruviana* and *gigas* corresponded to the northern and southern species, respectively, the properties of his data should be consistent with trait distributions from our larger dataset of measurements from these two species. To test this, we randomly subsampled our data without replacement to match Hellmayr’s sample sizes (i.e., we subsampled to *n*=8 for Southern males and *n*=4 for Southern females to match Hellmayr’s *gigas* sampling, and to *n*=25 for Northern males and *n*=8 for Northern females to match Hellmayr’s *peruviana* sampling), and generated 10,000 random datasets. For each of the 10,000 resampled datasets, we calculated trait means and coefficients of variation (CVs), generating expected distributions. We did this separately for each sex and species. We further calculated the magnitudes of trait difference between the two giant hummingbird species, generating expected distributions for differences in trait means. We then calculated the probability that Hellmayr’s data were compatible with each expected distribution, as would be expected if specimens had been selected without bias from individuals of the two species.

## RESULTS

### A neotype for *Patagona gigas* (now the Southern Giant Hummingbird)

To fix the identity of the unique species now known as the Southern Giant Hummingbird *Patagona gigas* (*sensu* Williamson et al. 2024), we designate as its neotype MSB:Bird:56142; CHILE: Valparaíso Region, locality Algarrobo, 13°20.916’ S, 71°38.294’ W, elevation 10 meters, collected 9 December 2018. Sex: female. Parts saved: partial skeleton, lung, heart, muscle, liver, eye, brain, and blood. Prepared by Selina M. Bauernfeind; personal catalog number SMB 516. Geolocator ID: BJ104, corresponding to annual migration tracks (Williamson et al. 2024). Specimen record: https://arctos.database.museum/guid/MSB:Bird:56142 (Fig. 2).

**Figure 2.**
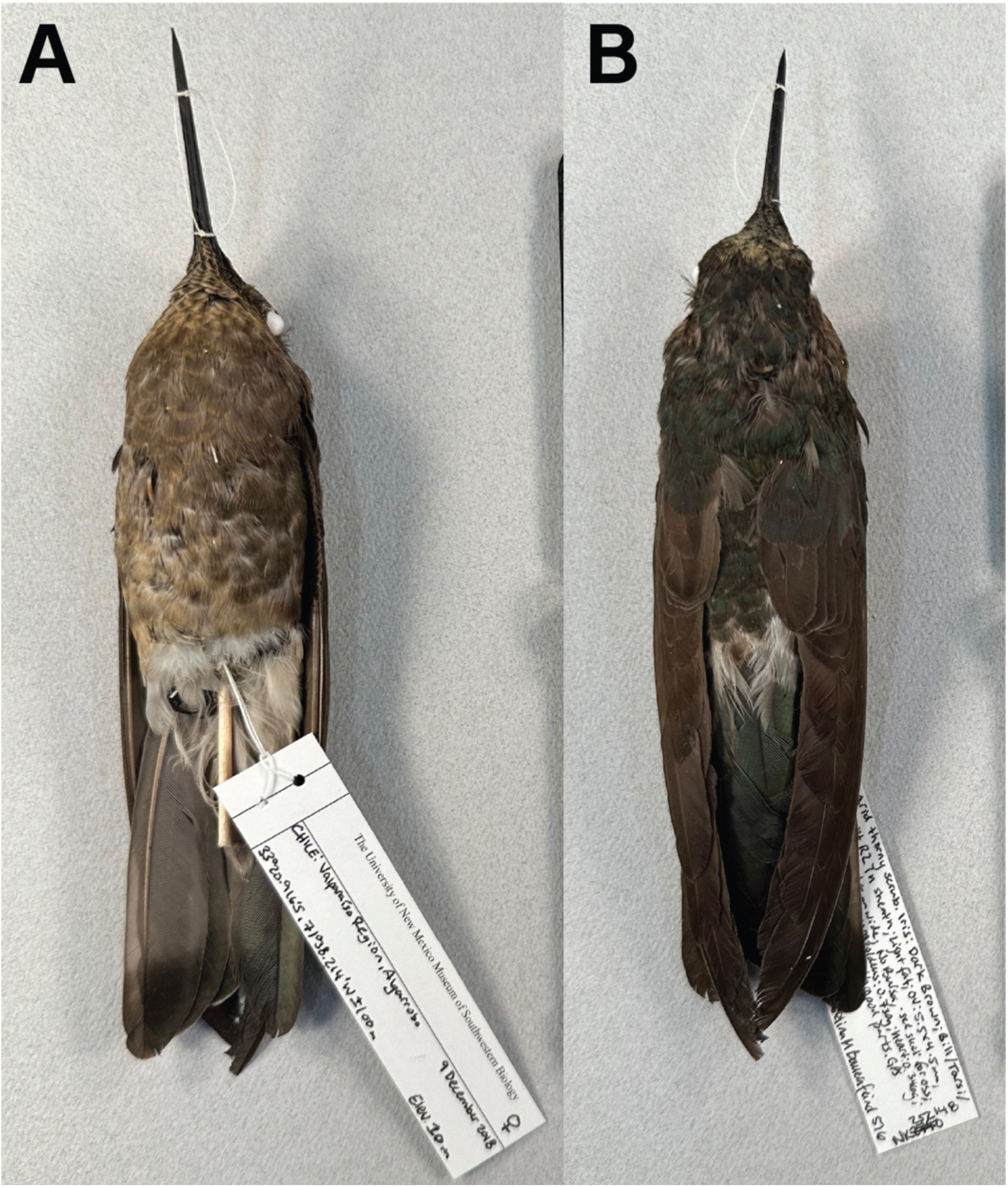
The *Patagona gigas* neotype, MSB:Bird:56142, housed at the Museum of Southwestern Biology at the University of New Mexico. The specimen, an adult female, was collected in Algarrobo, Valparaíso Región, Chile (13°20.916’ S, 71°38.294’ W) on 9 December 2018. Female. This specimen was unequivocally identified as a Southern Giant Hummingbird based on definitive nuclear and mitochondrial genomic, physiological, morphological, and migration tracking data (Williamson et al. 2024). **A** Ventral view. **B** Dorsal view. Specimen record and details are available from https://arctos.database.museum/guid/MSB:Bird:56142. Photos: [AUTHOR].

Why designate a neotype for *Patagona gigas*, rather than a lectotype from the putative syntype series? Art. 72.2.1 states that the type series consists of “…all the specimens on which the author established a nominal species-group taxon…in the absence of holotype designation, or the designation of syntypes, or the subsequent designation of a lectotype, all are syntypes and collectively they constitute the name-bearing type” (ICZN 1999). For *P. gigas*, the putative syntypes that could be considered candidates for lectotype designation may include: (1) Oudart’s plate of a “young male” described by Vieillot (1824), the specimen for which could not be located (Fig. S2); (2) the putative female voucher NHMUK-1933.11.14.36 (catalogued by NHMUK as possibly being Vieillot’s “adult female”; Fig. S3); and (3) the voucher MNHN-ZO-2023-396 (which Boucard believed might be “Vieillot’s type”; Fig. S4).

There are several reasons to doubt that either of the vouchers NHMUK-1933.11.14.36 and MNHN-ZO-2023-396 is a syntype of *P. gigas*. First, neither specimen possesses original tags––it is therefore impossible to confirm that they were used for Vieillot’s (1824) original description. It remains unknown whether NHMUK-1933.11.14.36 was the specimen originally acquired by Bécouer and subsequently examined by Vieillot in 1824; and there is no evidence that Vieillot ever examined MNHN-ZO-2023-396 (the latter specimen’s history and data are misaligned with Vieillot’s original description: for example, MNHN-ZO-2023-396 is sex unknown, yet Vieillot described an adult female; additionally, the MNHN does not recognize this specimen as a syntype). Second, neither voucher has a date of collection. NHMUK-1933.11.14.36 was obtained in 1933 after the purchase of George Loddiges’ collection; Loddiges, in turn, acquired the specimen from Jean-Baptiste Bécouer, who died in 1777, giving NHMUK-1933.11.14.36 an estimated but unconfirmed collection date prior to 1777. The MNHN-ZO-2023-396 specimen tags list no collection date for the skin, but the specimen is catalogued with a date of “avant 1842” (“before 1842”); however, Vieillot described *Patagona gigas* 18 years earlier (in 1824), and it is not possible to confirm that MNHN-ZO-2023-396 was collected prior to this date. Third, both vouchers have completely unknown provenance. There is no evidence that Bécouer ever traveled to Chile (or South America) to acquire NHMUK-1933.11.14.36, nor any specimen in his collection (Rookmaaker et al. 2006). The locality of MNHN-ZO-2023-396 is given only as “South America”. Hellmayr’s (1932) restriction of the type locality of *P. gigas* to “Valparaíso, Chile” was erroneous; it was based on the resemblance that he recognized between Oudart’s plate of a juvenile putative male and the immature female specimen FMNH-61676, collected September 23, 1924 from Caldera, Atacama Region, Chile, which is ∼830 km north of Valparaíso, Chile. Neither Oudart’s plate nor FMNH-61676 have any known link to NHMUK-1933.11.14.36 or MNHN-ZO-2023-396 (and there is no evidence that Vieillot’s “young male” and “adult female” syntypes were collected at the same time or from the same place). It is therefore not possible to confirm the identities of the putative syntype vouchers, and thus, it is not possible to confirm that voucher syntypes are extant.

Oudart’s plate of the “young male” is clearly indicated as part of Vieillot’s syntype series (Fig. S2). (Despite searching, we were unable to locate the voucher specimen of the individual depicted in the plate). Art. 74.4 of the Code states, “designation of an illustration or description of a syntype as a lectotype is to be treated as designation of the specimen illustrated or described; the fact that the specimen no longer exists or cannot be traced does not of itself invalidate the designation” (ICZN 1999). Oudart’s plate of the “young male” could therefore be considered an extant syntype and therefore a valid choice for a lectotype of *P. gigas*. However, Oudart’s plate depicts a juvenile and it is established that the Southern and Northern Giant Hummingbird juveniles cannot be reliably distinguished by plumage (Williamson et al. 2024). Furthermore, the plate, which Hellmayr (1932) noted was “…none too good…”, does not illustrate any diagnostic phenotypic characteristics of the Southern Giant Hummingbird, the species which it is meant to denote. The locality, provenance, and date of collection of the specimen voucher illustrated in the plate are additionally unknown, so it cannot be verified whether it truly corresponds to the Southern Giant Hummingbird. The collector of the specimen is also unknown. Thus, Oudart’s “young male” (sex unconfirmed) plate violates Recommendations 74C (and by extension, 73C) and 74E and is therefore unsuitable as a lectotype for *P. gigas.* Art. 72.2 states, “…If no name-bearing type is believed to be extant a neotype may be fixed…” (ICZN 1999, 2003).

In accordance with Art. 75.3 of the Code, there is an exceptional need to designate a neotype for *P. gigas*. The designation of neotype specimen MSB:Bird:56142 satisfies the full list of qualifying conditions of the Code: This designation is made with the express purpose of clarifying both the taxonomic status *and* the type locality of the nominal taxon *P. gigas* (Art. 75.3.1); the neotype specimen MSB:Bird:56142 is unequivocally identified as a Southern Giant Hummingbird based on definitive nuclear and mitochondrial genomic, physiological, morphological, and migration tracking data; these features clearly distinguish MSB:Bird:56142 from individuals of the Northern Giant Hummingbird (Williamson et al. 2024; Art. 75.3.2); the archived specimen data and above neotype description are sufficient to ensure recognition of the neotype, MSB:Bird:56142 (see Fig. 2; Art. 75.3.3); no name-bearing types are believed to be extant for the aforementioned reasons (Art. 75.3.4); the neotype MSB:Bird:56142 has phenotypic characteristics consistent with what is known of the name-bearing syntypes originally described by Vieillot (1824) and subsequently described by other authorities; the neotype MSB:Bird:56142 is additionally a female, matching Vieillot’s original description of one of the syntypes (Art. 75.3.5); the neotype MSB:Bird:56142 was collected in Valparaíso Region, Chile, the type locality that Hellmayr (1932) intended to designate, but which he had restricted in error (Art. 75.3.6); and, the neotype MSB:Bird:56142 is housed at the Museum of Southwestern Biology at the University of New Mexico, a recognized scientific and educational institution that maintains a research collection and which possesses proper facilities for preserving name-bearing types that are accessible for study (Art. 75.3.7). This neotype designation is in accordance with Recommendation 75A. [The authors have sought feedback from the Working Group on Avian Nomenclature to satisfy Recommendation 75B and will include the appropriate ZooBank LSID after Working Group input] (ICZN 1999, 2003).

As stated in Art. 76A.2, “A statement of a type locality that is found to be erroneous should be corrected”. Art. 76.3 states, “The place of origin of the neotype becomes the type locality of the nominal species-group taxon, despite any previously published statement of the type locality”. Thus, the type locality for the Southern Giant Hummingbird (*Patagona gigas*) should henceforth be considered Algarrobo, Valparaíso Region, Chile, the collection locality of the designated neotype, which corrects the previous erroneous type locality restriction. The designation of MSB:Bird:56142 as the neotype fixes the identity of *Patagona gigas*, ensuring nomenclatural stability (ICZN 1999, Jiménez-Mejías et al. 2024).

### How an unknown migratory route led to a muddied taxonomy

Numerous published accounts, both historical and modern, have mentioned giant hummingbird migration or presence/absence of *Patagona* at certain times of the year in Chile with only sparse details (e.g., Gould 1861, Barros 1952, Aguirre 2001, González-Gómez and Valdivia 2005, Estades et al. 2008, Speziale and Lambertucci 2009, Velásquez-Noriega et al. 2021, Medel et al. 2022, Rueda-Uribe et al. 2023). One publication stated that *Patagona* migrates 2,000 km (Rodríguez-Flores et al. 2019), but no information about the migratory route or sources for distance traveled were provided. Two previous studies used coarse-scale citizen science data to estimate the magnitude of elevational shift experienced seasonally by migratory populations (Williamson and Witt 2021a, Rueda-Uribe et al. 2023); both generated what we now know are underestimates.

Lack of information about the migration of southern populations has led to persistent uncertainty about the ranges of each giant hummingbird form. Throughout history, the giant hummingbird range has generally been described as encompassing the Andes, but texts are not consistent and have often omitted mention of Argentina (e.g., Gould 1861, Boucard 1893). In particular, the range limits of *peruviana* had never been described correctly. Hellmayr (1932) wrote that “*P. g. peruviana* … replaces the typical form in the Andes of Ecuador, Peru, Bolivia, and extreme northern Chile. It is probably also this race that nests in northwestern Argentina, although I have not been able to examine undoubted breeding specimens” (Hellmayr 1932). In 1945, Peters wrote, “The breeding bird of northwestern Argentina (Jujuy, Catamarca, and Tucumán) is probably referable to [*P. g. peruviana*]” (Peters 1945). In 1952, Zimmer identified several Southern Giant hummingbirds from Argentina and Bolivia as *peruviana* (Zimmer 1952). The belief that the race *peruviana* occurs in northwest Argentina has been supported by many key historical accounts (Barros 1952, Zimmer 1952, Ruschí 1961, Koepcke 1983, Fjeldså and Krabbe 1990) and has continued to persist into recent years (e.g., Sánchez Osés 2003, Pearman and Areta 2020, Velásquez-Noriega et al. 2021). More recently, in *Birds of Chile: A Photo Guide*, Howell and Schmitt (2018) stated that there is “no range overlap” between *peruviana* and *gigas* (Howell and Schmitt 2018). However, we now know that northern Argentina contains only the Southern Giant Hummingbird, and that the two species overlap substantially in range during the non-breeding season.

Uncertainty about migration and confusion about the numbers of possible subspecies/forms, and their respective distributions (e.g., Fjeldsa and Barbosa 1983), has additionally resulted in misleading generalizations about elevations of residence. Woods (1998) suggested that the two forms, *peruviana* and *gigas*, might be elevational replacements and incompletely characterized the Southern Giant Hummingbird range, writing, “The northern subspecies of Giant Hummingbird *P. g. peruviana* regularly reaches paramo and puna habitats in Ecuador and Peru at 3,500-4,000 m, while the southern *P. g. gigas* occurs seasonally to below 2,000 m in central Chile and adjacent Argentina” (Woods et al. 1998). Ruschi (1961) stated that *P. g. peruviana* is found in the Andes, varying in altitude from 2,500 to 5,000 meters (Ruschí 1961) but he did not mention *P. g. gigas* or its range. In many other texts, it is not possible to determine which form of giant hummingbird is being discussed.

We now know that the giant hummingbirds are comprised of two phenotypically cryptic species that overlap at the same sites and elevations during the southern hemisphere winter, approximately May–September (Williamson et al. 2024). The Southern Giant Hummingbird is a long-distance latitudinal migrant, traveling up to 8,300 km roundtrip in a circular loop across the Andes (Williamson et al. 2024). At least some (i.e., Chilean) Southern Giant Hummingbird populations shift >4,100 meters in elevation twice annually, making this species one of ∼100 taxa that undertake elevational niche-shift migration (ENSM), or physiologically-taxing ‘extreme’ shifts in elevation between the breeding and non-breeding seasons (Williamson and Witt 2021a, b; Williamson et al. 2024, Ivy and Williamson 2024). Although Southern Giant Hummingbird populations from Argentina also spend the non-breeding season at high elevations in central Peru, their migratory routes and degrees of elevational shift have not yet been studied. Resolution of longstanding mysteries about giant hummingbird migration was made possible with the use of miniaturized tracking devices and genomic sequencing—technologies that were not available to Darwin, Johow, or the other workers cited above—that has finally permitted clarification of taxonomy within this clade.

### The giant hummingbirds are not “easily” distinguished

Contrary to Hellmayr’s (1932) findings, our measurements of >600 specimens revealed substantial overlap between species for bill, wing, and tail lengths within each sex (Fig. 3). Hellmayr’s *peruviana* measurements were significantly larger than our Northern Giant Hummingbird measurements in sex-specific comparisons of wing and tail, respectively, and female bill length (Fig. 3b–f). For each sex, Hellmayr’s data also showed significant differences between *gigas* and *peruviana* tail measurements, whereas we found none (Fig. 3c,f). When we conducted a Principal Components Analysis (PCA) using bill, wing, and tail measurements separately for each sex, Hellmayr’s data clustered tightly and were shifted to the outer edges of the expected distributions (Fig. 4). In this same analysis, the MNHN male *peruviana* lectotype (MNHN-ZO-MO-1989-340), an apparent Northern Giant Hummingbird, and the *boliviana* holotype (MNHN-ZO-MO-1989-342), a Southern Giant Hummingbird of unknown sex, both fell within Hellmayr’s range for *peruviana*, and within a morphological zone of overlap between male Northern and Southern Giant Hummingbirds; when analyzed with female data, MNHN-ZO-MO-1989-342 was an outlier for both species (Fig. 4). The MNHN *peruviana* female paralectotype (MNHN-ZO-MO-1989-341) fell more closely in line with Southern Giant Hummingbird measurements, but its identity was not clearly identifiable based on measurements (Fig. 4).

**Figure 3.**
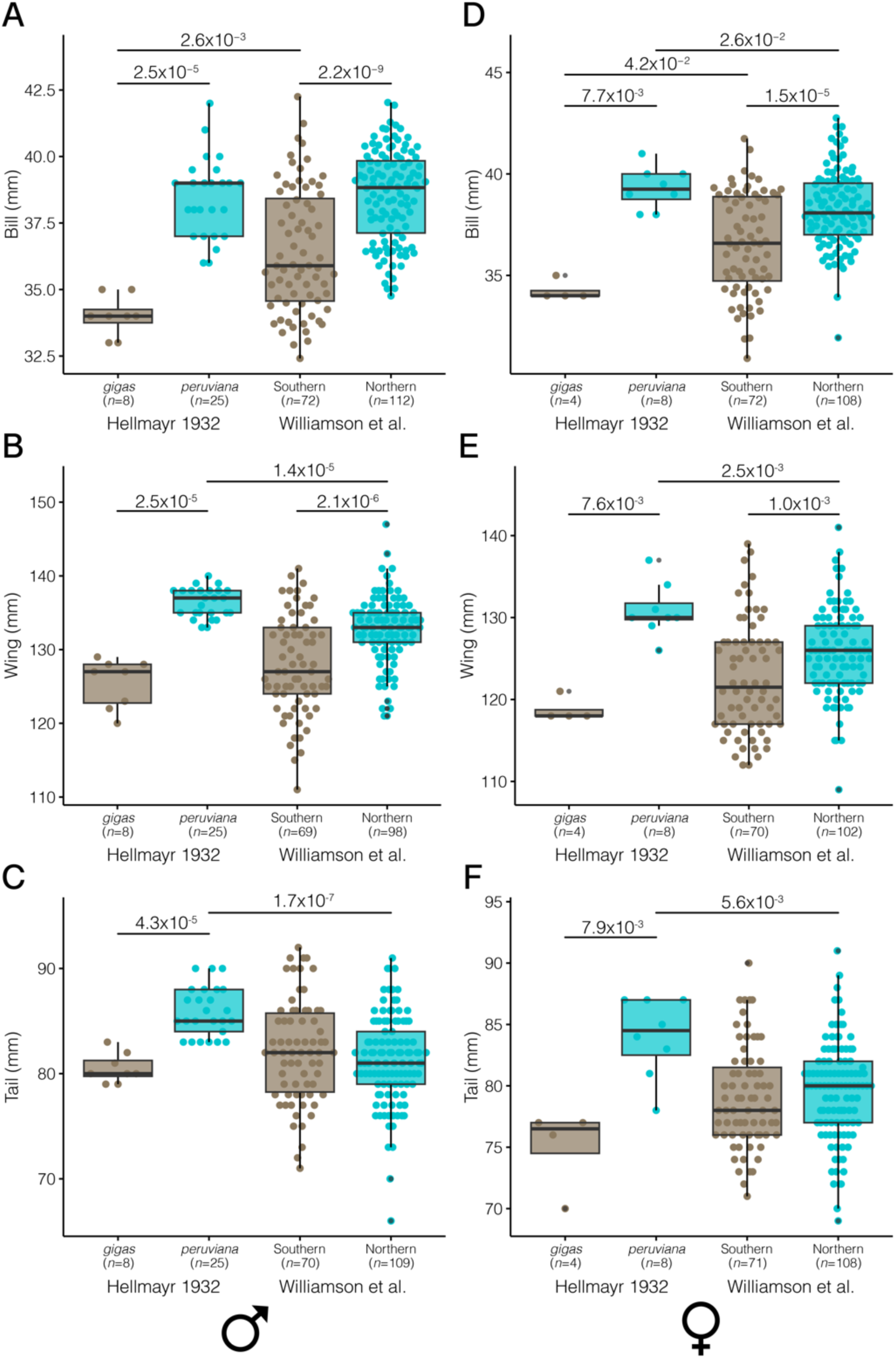
Comparison of morphological data (bill length, wing chord, and tail length; all in mm) between Hellmayr (1932) and Williamson et al. (present study). **A–C** Males. **D–F** Females. Hellmayr compared adults of the two suspected forms, *gigas* and *peruviana* (*n*=45 specimens total). Williamson et al. compared adults of the two species, the Southern and Northern Giant Hummingbird (*n*=448 individuals total). Sample sizes for each comparison are reported on the x-axis and significant *p*-values are indicated above each comparison (from t-tests or non-parametric Wilcoxon Rank Sum tests). Box plot horizontal lines represent median values. Each point is a value from a single individual.

**Figure 4.**
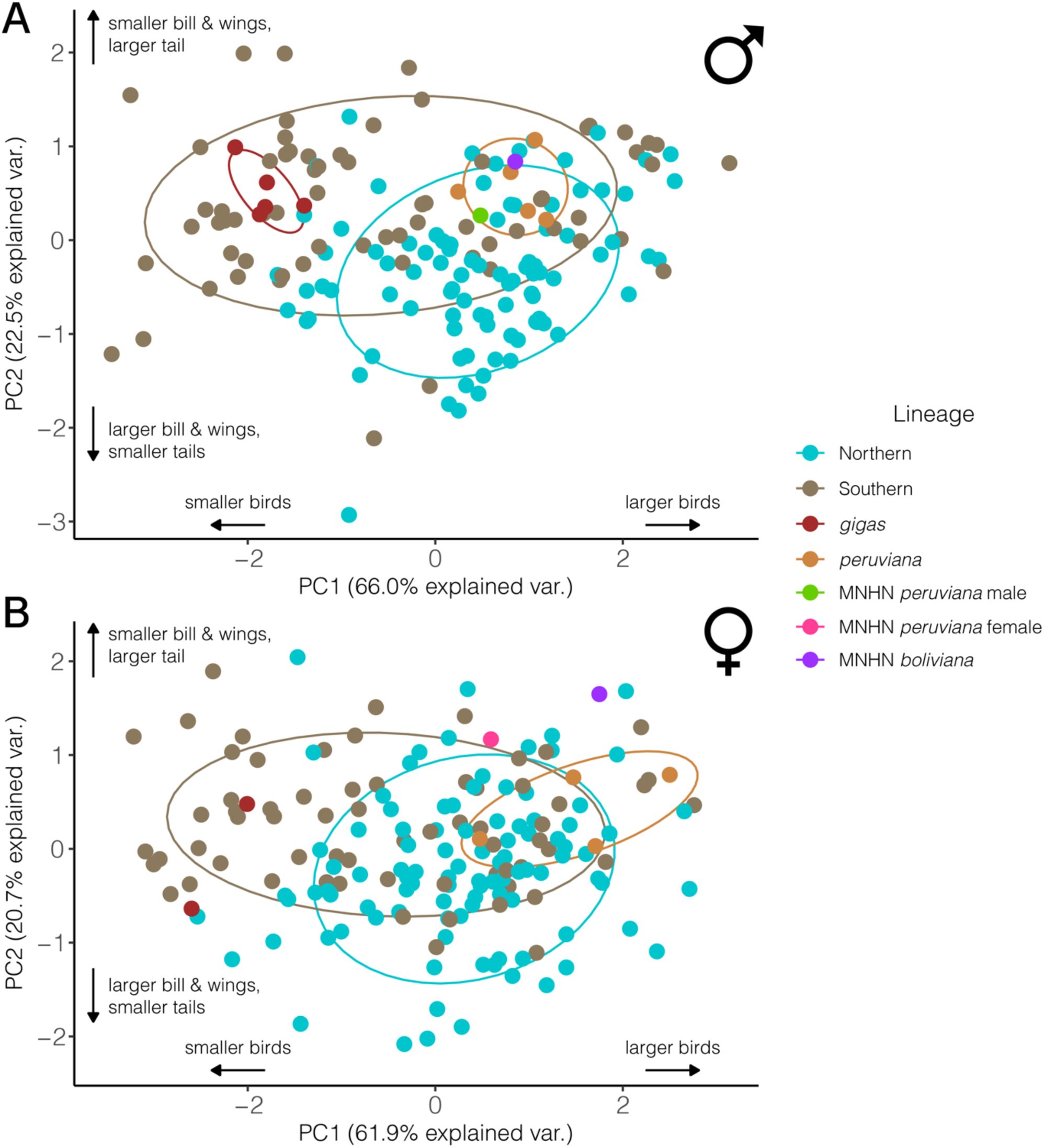
Principal Component Analysis (PCA) of bill, wing, and tail measurements for *gigas* and *peruviana* individuals from Hellmayr (1932) and Southern and Northern Giant Hummingbirds from Williamson et al. (present study) with MNHN *peruviana* specimens and the *boliviana* holotype. Hellmayr’s data were entered numerically, rather than by individual; we have therefore included only records with single individuals per site. **A** Males (*n*=172 individuals); **B** Females (*n*=174 individuals). The male *peruviana* lectotype (MNHN-ZO-MO-1989-340), female *peruviana* paralectotype (MNHN-ZO-MO-1989-341), and *boliviana* holotype (MNHN-ZO-MO-1989-342) are plotted for reference; the *boliviana* holotype is sex unknown and is thus included in both PCAs. The female *peruviana* specimen is plotted despite the tip of its maxilla being broken (Patrick Bousses, *pers. comm.*); however, *Patagona* maxilla and mandible lengths are typically equal. For reference, bill measurements: *peruviana* male: 35.7 mm; *peruviana* female 35.8 mm (mandible only); *boliviana*: 35.8 mm.

Hellmayr’s differences in trait means between forms were noticeably larger, and CVs in his dataset noticeably smaller, than in the expected distributions generated from 10,000 resampled datasets (Figs. S5–S6, Tables S2–S4). For all sex-trait combinations, his CVs and differences in trait means were statistically improbable and unrealistically small (Figs. S5–S6; Supplement). For example, the difference between Hellmayr’s *peruviana* and *gigas* mean wing measurements was ∼2x greater for males and ∼4x greater for females than the respective differences we found between Northern and Southern Giant Hummingbirds (Fig. 3).

### Misidentification of specimens in historical texts and museum collections

Many historical texts do not provide specimen identifiers but, in several instances, it is possible to diagnose specimen identities using descriptive information provided. Hellmayr (1932) writes of three specimens that “…are in every particular typical of the large northern form…an adult male from Tilcara, Jujuy (Nov. 24, 1905; L. Dinelli), and a couple from Lara, Tucumán (Feb. 12, 1903; G. A. Baer), all in the Tring Museum” (Hellmayr 1932). These specimens were formerly at NHMUK but were moved to the AMNH in 1932 after Lord Rothschild sold his collection (Lanyon 1995, Roselaar 2003; Mark Adams *pers. comm.*; Tom Trombone, *pers. comm.*) The adult male from Tilcara described by Hellmayr is AMNH-479595, definitively identified as a Southern Giant Hummingbird based on targeted sequence capture of ultra-conserved elements (UCEs), known breeding season collection locality, and diagnostic plumage characteristics (Williamson et al. 2024). The other two Lara, Tucumán individuals mentioned are AMNH-479593 and AMNH-479594; both are juvenile Southern Giant Hummingbirds, for which Williamson et al. (2024) also provide UCE data. The fact that Hellmayr believed these individuals from Argentina were “in every particular typical” of *peruviana*, when in fact, they are Southern Giant Hummingbirds ranging in age from juveniles to adults highlights the difficulty of distinguishing the two forms by plumage.

Due to these challenges, many historic and modern Southern Giant Hummingbirds (*gigas*) were misidentified as Northern Giant Hummingbirds (*peruviana*). The reverse does not seem to be true: We have found no Northern Giant Hummingbirds misidentified as Southern Giant Hummingbirds. At least eleven major museum collections whose specimens we have examined possess giant hummingbird specimens from each Peru, Bolivia, Argentina, and Chile that are catalogued or labeled as *P. g. peruviana*, and which contain a mix of both the Northern and Southern Giant Hummingbird. Of the 603 total adult and juvenile specimens with locality data whose catalog records and tags we analyzed, only 269 individuals (44.6%) are catalogued with subspecific identifiers. A total of 74 individuals (32.74% of specimens) labeled “*peruviana*” are actually Southern Giant Hummingbirds (e.g., AMNH-139179, ANSP-120834, CM-52921, CUMV-52570, FMNH-67518, LSUMZ-37404, MCZ-262196, USNM-529148; Fig. 5; Table S5). Two individuals (ANSP-145417 and CM-94400) from Bolivia were labeled with both “*peruviana*” and “*gigas*” (on the tag of ANSP-145417, pencil marks are visible where “*peruviana*” was erased and “*gigas*” was written; this individual still bears “*peruviana*” on the back of its tag).

**Figure 5.**
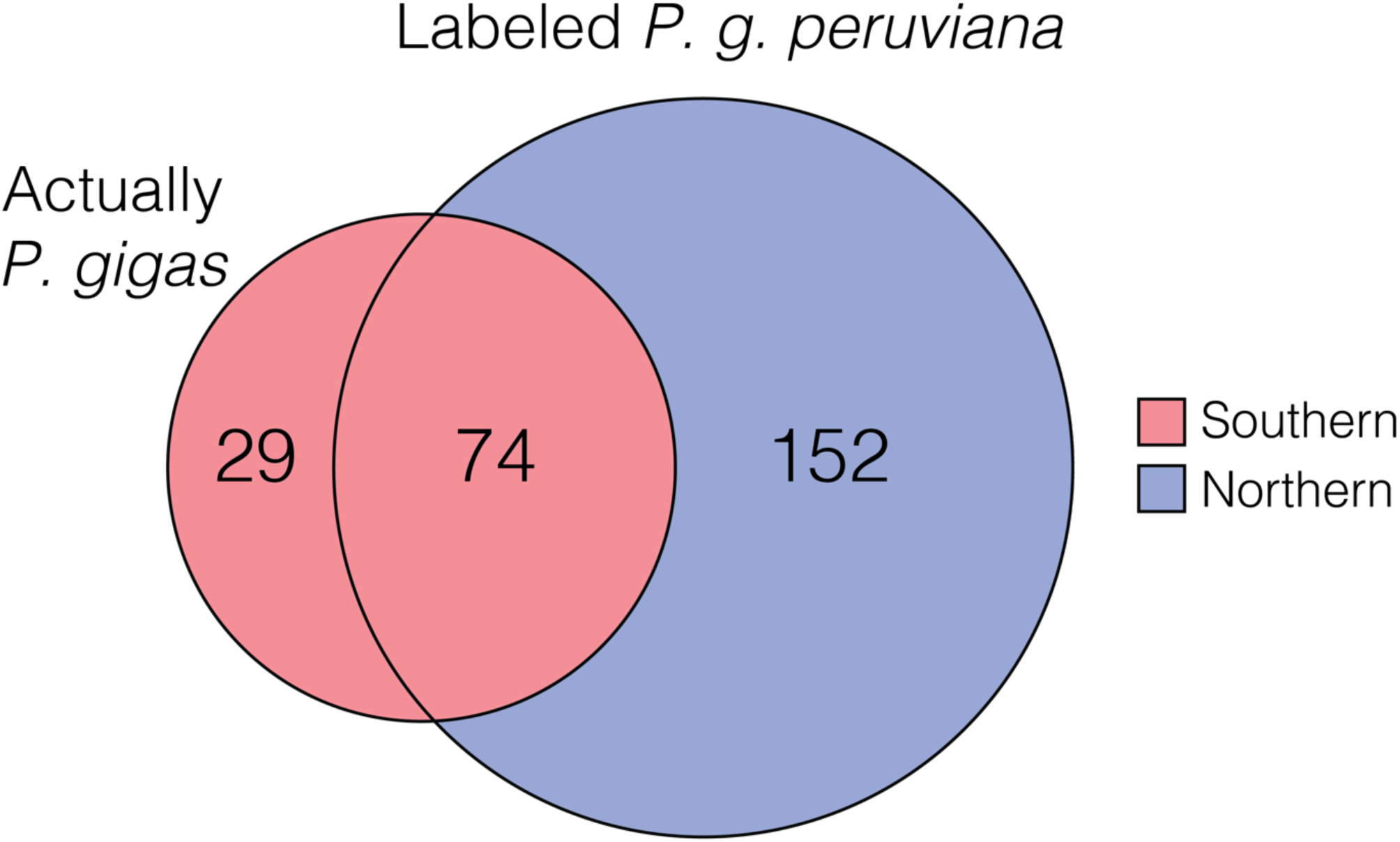
Visual representation of the *Patagona* specimen record as it pertains to previous versus current identifications (see Table S5 for full list). Of the 603 total adult and juvenile specimens with locality data whose catalog records and tags we analyzed from eleven major museum collections, a total of 269 individuals (∼44%) are catalogued with subspecific identifiers. Of these, 29 specimens are labeled as *P. gigas* and pertain to the Southern Giant Hummingbird (*Patagona gigas*, sensu Williamson et al. 2024). A total of 226 specimens with known lineage are labeled as *P. g. peruviana*; of these, 152 specimens are correctly identified as Northern Giant Hummingbirds (*P. peruviana*) and 74 specimens are incorrectly identified as *P. g. peruviana* and are Southern Giant Hummingbirds (*P. gigas*).

Typically, any individual collected in Ecuador, Peru, Bolivia, and northwestern Argentina that appeared large and had reddish underparts was assigned to *peruviana* without substantial scrutiny; by this measure, many rufous-toned *gigas* individuals, especially those that seemed larger, would appear to correspond to *peruviana* (e.g., Fig. 1). The name *peruviana* has thus been applied to specimens on the basis of oversimplified morphology, plumage, and assumptions about range. However, the extent to which *gigas* specimens have been misidentified (and thus, the name *peruviana* misapplied) in global collections is difficult to ascertain because the vast majority of catalogued specimens (55.4%) do not have subspecific identifiers.

### Nomenclatural recommendations

The neotype designated here (MSB:Bird:56142) corresponds unambiguously to the Southern Giant Hummingbird, and we thus recommend continued use of *Patagona gigas* for this newly-recognized species. The name *peruviana* has a contentious history: Although it has always intended to apply to the ‘larger, northern form’ of the giant hummingbird, there has been longstanding uncertainty about how the ‘larger, northern form’ is defined, and about how the name was applied to individuals of certain characteristics. Consequently, the name *peruviana* has been applied, variably, to the two giant hummingbird species based on a combination of erroneous or incomplete information about plumage, morphology, range, and seasonal occurrence. The *peruviana* lectotype (MNHN-ZO-MO-1989-340) appears to correspond in plumage to the Northern Giant Hummingbird. Contrary to Areta et al. (2024)’s statement that MNHN-ZO-MO-1989-340 is “unambiguously” identifiable as the northern taxon, there is reasonable room to doubt this plumage-only-based assessment and the identity of the lectotype remains unconfirmed without genetic sequencing. However, following rules set forth by the Code, the name *peruviana*, the current senior synonym, should be adopted for the Northern Giant Hummingbird.

## DISCUSSION

### Historical oversimplification of complexity within and between *Patagona* species

Generations of ornithologists have pondered whether the giant hummingbirds comprise one or more forms, and have attempted to characterize their distinguishing criteria. In his 1932 text, Hellmayr concluded that there were strong morphological differences “clearly” indicating the existence of two forms. However, the analysis by Williamson et al. (2024), as well as our present comparison, which included data from an additional 248 adults from across the range, revealed substantial overlap in sex-specific mensural traits between Northern and Southern Giant hummingbirds. Hellmayr’s data did not show this overlap between forms, even when controlling for sex (Fig. 3). Areta et al. (2024)’s assertion that Hellmayr’s results were “clearly mirrored” by Williamson et al. (2024) is an overstatement. The *peruviana* measurements in Hellmayr’s dataset were significantly greater than those in our dataset for all comparisons except male bill length (Fig. 3). When compared with 10,000 resampled trait distributions, Hellmayr’s results were highly improbable, suggesting that the birds were identified, at least in part, based on their measurements and incorrect assumptions about inter-taxon phenotypic variation. The differences between taxa in Hellmayr’s data are too large and the variability within taxa too small to have been produced by an unbiased sampling of the two species. In this way, Hellmayr’s measurements do not comprise valid taxonomic evidence.

Why did Hellmayr’s (1932) data appear so decisive in distinguishing two *Patagona* taxa? One explanation is that Hellmayr included misidentified specimens in his dataset; we know this occurred because at least several *P. gigas* specimens were included in his “*peruviana*” series (e.g., AMNH-479593, AMNH-479594, and AMNH-479595). Including misidentified individuals would diminish between-taxon differences and inflate variability within taxa––the opposite of the pattern observed in Hellmayr’s data. Alternatively, Hellmayr may have used a combination of plumage assessment, measurements, and locality to assign each specimen to a taxonomic category; thus, he may have started with the *a priori* assumption that the two taxa were non-overlapping in size traits and range, and may then have included or omitted data that fit with this hunch. Considering the magnitude of the challenge inherent in identifying giant hummingbirds based on plumage, it may have been Hellmayr’s best option to make these educated guesses, seeking consilience of evidence as he tried to fit a taxonomy to this difficult group. While this approach may have been reasonable given the tools of the era, the resulting conclusions were therefore based on overly-generous distinctions that consequently led to further misidentification. Hellmayr is not the only worker to have studied the giant hummingbirds; however, his 1932 text is widely regarded as definitive for giant hummingbird identification and it is perhaps the only known text to provide raw data, permitting present-day re-analysis.

Robust analysis of morphology using linear discriminant analysis, principal component analysis, and comparisons of means on our dataset spanning the entire *Patagona* distribution has reinforced that individuals of each the Northern and Southern species may be difficult to distinguish even with a suite of measurements used in concert (Figs. 3–4). It is therefore unwise to characterize the two giant hummingbird species on the basis of any single morphological trait, and it would be imprudent to use any single measurement to attempt to identify individuals to species, as was done recently by Areta et al. (2024).

One underappreciated component of giant hummingbird identification is the extent of individual plumage variation that exists *within* each species (Fig. 1). The authors’ study of >600 wild-caught individuals and museum specimens suggests that the extent of cinnamon coloration appears to be related to age; even so, each species possesses a high degree of plumage variation that appears unrelated to sex. Hellmayr (1932) and Zimmer (1952) each noted this variation, suggesting that it might be random; Simon (1921) noted age-associations with plumage color corroborated by our study. Even among adults, the range of variation in the throat color and pattern of Southern Giant Hummingbirds overlaps that of Northern Giant Hummingbirds. Although it is generally possible to distinguish the two species using plumage characteristics, because within-species variation in throat color and pattern may overlap, many individuals are best left unidentified when observed during locations and seasons of co-occurrence. A clearer understanding of the seasonal elevational and latitudinal ranges of both species will be essential to field identification and further understanding migration, seasonal zones of overlap, and regions where the two species hybridize.

### Distribution and identification of the giant hummingbirds

Here, we provide details about Northern and Southern Giant Hummingbird identification, seasonal distributions, elevational ranges relevant to ornithologists and birders.

#### Northern Giant Hummingbird (Patagona peruviana)

**Plumage:** Adult Northern Giant Hummingbirds typically have whitish or buffy-colored throats with dark blackish streaks (but note that throat color is variable, from very high to more subtle contrast), a subtle cinnamon patch at the base of the throat (giving the throat base a more rufous color), and a whitish eye-ring with post-ocular spot (Fig. 1). Wings and bill average longer, but identification by measurements alone is often not possible.

**Distribution and range:** The Northern Giant Hummingbird is found year-round in extreme southern Colombia, Ecuador, Peru and northern Chile (Arica and Parinacota Region); it is not yet confirmed to occur in Bolivia, and there are no records (photographs, recordings, or specimens) of this species in Argentina. It spans an elevational range of ∼1,800-3,600 m in Ecuador, ∼1,900-4,300 in Peru, and ∼1,900-3,800 m in Chile. In Peru, observers should carefully scrutinize giant hummingbirds during periods of seasonal overlap (∼May–Sept), as the two *Patagona* species co-occur in many of the same sites and elevations. Individuals observed throughout Peru from November–April are most likely Northern Giant Hummingbirds; individuals observed in central to southern Peru from May–September could plausibly be Northern or Southern Giant Hummingbirds. The Northern Giant Hummingbird may appear very similar in plumage to the Southern Giant Hummingbird (Fig. 1), particularly in poor lighting conditions or when seen in flight. This species is found in arid and semi-arid habitats on the west slope of the Andes and in intermontane valleys, including montane scrub, hedgerows, agricultural areas, open woods (including *Polylepis*), and gardens, often with *Agave*, *Puya*, and Cactaceae spp. Across its range, it tends to associate with fast-flowing streams.

#### Southern Giant Hummingbird (Patagona gigas)

**Plumage:** The Southern Giant Hummingbird tends to have a warm brown to cinnamon throat with subtly contrasting grayish-green streaks (but note that throat contrast is variable and may approach Northern in coloration), occasionally with a subtle cinnamon patch at the base; some individuals may have extensive rufous throughout the throat and breast. The eye-ring and post-ocular spot are subtle (buffy-colored, if present) to absent, giving Southern individuals a ‘blank-faced’ appearance (Fig. 1). Wings and bill average shorter.

**Distribution and range:** The Southern Giant Hummingbird is found in central Chile, central through northwest Argentina, Bolivia, and Peru (to central Ancash Department). This species breeds from sea level to ∼2,000 m in Chile, above ∼2,500 m in Argentina, and ∼2,100–4,150 m in Bolivia. After breeding, Southern Giant Hummingbirds in central Chile migrate (beginning ∼mid-February to March) to Peru, passing through the breeding grounds of populations from northwest Argentina. At least some populations that breed in northwest Argentina also migrate to central Peru, but details about migration are as yet unknown. In Peru, the Southern Giant Hummingbird is found on the nonbreeding grounds, up to 4,300 m, at the same sites and elevations as the Northern Giant Hummingbird. Southern Giant hummingbirds are estimated to comprise ∼20–40% of the nonbreeding population in Peru during May–September (Williamson et al. 2024); however, further work is needed to refine our understanding of this species migratory stopover sites and nonbreeding range. Individuals observed in Bolivia should be carefully evaluated, particularly in the north, to rule out presence of the Northern Giant Hummingbird. Individuals in Chile and Argentina may overwinter. Observers should carefully scrutinize individuals in northern Chile (Arica and Parinacota Region), particularly during ∼August–October when Southern Giant Hummingbirds migrate through the region (and may pass through the range of Northern Giant Hummingbirds).

#### Juvenile giant hummingbirds

The plumage of juvenile giant hummingbirds (called juvenal plumage) of both species is nearly identical (Fig. 1); previous workers have noted the difficulty of distinguishing juveniles (e.g., Zimmer 1952, Ortiz-Crespo 1974). The juveniles of both species have deep cinnamon throats, breasts, and bellies that fade to duller brown as birds molt into immature plumage; emerald-toned heads, backs, and wings with buffy-tips to contour feathers; a pronounced white rump patch that may extent mid-way up the back; and fresh white tips to primaries and rectrices (Mulsant and Verreaux 1876; Fig. 1). Dark black streaking seems to appear sooner on the throats of Northern Giant Hummingbird juveniles; presence of this character may prove useful for identification. Season and locality should be carefully evaluated when observing juveniles.

### Conclusions

The migration-tracking, genomic, physiological, and morphological study conducted by Williamson et al. (2024) demonstrated that elevational migrant and high-elevation resident populations of the giant hummingbird comprise distinct species, now known as the Southern and Northern Giant Hummingbird, respectively (Williamson et al. 2024). Before the introduction of these novel data on migration and genetics, the taxonomy of giant hummingbirds was impossible to disentangle, in part because of erroneous assumptions about ranges and oversimplified identification criteria. The first described species in the genus, *Patagona gigas,* and the taxon *peruviana,* have had confusing histories. The putative voucher syntypes for *P. gigas* have unconfirmed identities, unknown collection dates, and dubious provenance; lacking critical information, they cannot be confirmed as extant. The final syntype, a plate depicting a juvenile putative male, does not possess any identifying features of the Southern Giant Hummingbird and is thus unsuitable as a lectotype. To clarify the status of the Southern Giant Hummingbird (*Patagona gigas*; *sensu* Williamson et al. 2024), we designate a neotype for *P. gigas* by selecting a modern specimen of known provenance linked to definitive nuclear and mitochondrial DNA sequence data, and for which annual migration data exist (MSB:Bird:56142); this designation additionally corrects the previous erroneous type locality restriction. The newly-designated *peruviana* lectotype appears to correspond to the Northern Giant Hummingbird in plumage, and the name *peruviana* therefore has nomenclatural precedence as a senior synonym. Thus, despite introduction of the name *chaski* concomitantly with definitive delimitation of species limits, phenotypic characterizations, physiological and behavioral differences, and year-round distributions for the two *Patagona* species (Williamson et al., 2024), *chaski* is a junior synonym of *peruviana*. We recommend DNA sequencing to confirm the identity of the *peruviana* lectotype.

The two cryptic giant hummingbird species are highly challenge to distinguish by plumage and morphology alone, as confirmed by our analysis of 608 wild-caught individuals and museum-vouchered natural history specimens spanning 1864–2023, including both species across the Andean range. These findings contrast the longstanding idea that the two forms were “easily” distinguished, additionally highlighting how inaccurate historical diagnoses led to misidentification in the field and in collections. We clarify the historical record by providing a list of specimens misidentified in museums. Finally, we offer detailed species accounts with notes on plumage diagnosis, and seasonal latitudinal and elevational ranges, to aid ornithologists and birders in further field study.

As ornithologist Ted Parker once noted, “A taxonomic puzzle illuminates everything else about a bird. The thing to keep in mind––because some people here and elsewhere think finding a new species is the ultimate thing––is how much we have learned about all of these birds” (Stap 1991, pg. 160).

## Acknowledgements

We are grateful to Frank Rheindt, Thomas Pape, Pamela Rasmussen, Oscar Johnson, Alex Figueroa, Glaucia Del-Rio, Shawn Billerman, Terry Chesser, and J. V. Remsen, Jr. for helpful discussion on taxonomy and nomenclature. Field assistance was provided by Kyana Montoya and Marlon Chagua, French language translation by Peter Truog, and distribution information by Leonardo Campagna. We are grateful to the museum curators, collections managers, staff, and students who facilitated this work: Mark Adams, Mike Andersen, Clare Atkinson, John Bates, Bentley Bird, Sharon Birks, Patrick Boussès, Serina Brady, Mariel Campbell, Eamon Corbett, Joel Cracraft, Charles Dardia, Carla Dove, Scott Edwards, Rob Faucett, Mary Margaret Ferraro, Jérôme Fuchs, John Gerwin, Gary Graves, Emily Griffith, Shannon Hackett, Andy Johnson, John Klicka, Andy Kratter, Terry Lott, Irby Lovette, Ben Marks, Nick Mason, Jon Merwin, Chris Milensky, Rob Moyle, Jose Antonio Otero, Nathan Rice, Mark Robbins, Sahid Robles-Bello, Vanya Rohwer, Sam Rutledge, Dave Steadman, Paul Sweet, Jeremiah Trimble, Tom Trombone, Hein van Grouw, Jason Weckstein, and J. V. Remsen, Jr. Collections access, data, and historical information were provided by the American Museum of Natural History, Academy of Natural Sciences of Drexel University, Carnegie Museum of Natural History, Centro de Ornitología y Biodiversidad, Cornell University Museum of Vertebrates, Field Museum of Natural History, Louisiana State University of Museum of Natural Science, Museum of Comparative Zoology at Harvard University, Muséum National d’Histoire Naturelle, Museum of Southwestern Biology, Natural History Museum at Tring, and the Smithsonian National Museum of Natural History. This manuscript is under review by the Working Group on Avian Nomenclature. Data from wild birds measured in Peru were collected under Servicio Nacional Forestal y de Fauna Silvestre RD-000056-2023-DGGSPFFS-DGSPFS. IACUC approval was granted under Cornell University protocol 2022-0127.

## Author contributions

JLW and CCW conceived the idea. All authors collected the data. JLW curated the data and obtained funding. JLW and CRG analyzed the data. JLW, CRG, and BWR made figures and tables. JLW, TV, and CCW contributed critical resources. JLW wrote the initial draft. JLW and CCW substantially revised the manuscript with input from all authors.

## Conflict of interest

The authors declare no conflicts of interest.

## Funding

This work was supported by an NSF Postdoctoral Research Fellowship in Biology (DBI-2208924) and a Cornell Lab of Ornithology Edward W. Rose Postdoctoral Fellowship to JLW. Collections visits were supported by an American Philosophical Society Franklin Research Grant.

## Data availability statement

Data from Williamson et al. (2024) analyzed here are available on Dryad: https://datadryad.org/stash/dataset/doi:10.5061/dryad.44j0zpcnp. Unpublished data will be uploaded to a repository upon manuscript acceptance.

## SUPPLEMENT

### The Whitely series of *Patagona* specimens

Henry H. Whitely, Jr., a British naturalist and explorer particularly interested in ornithology (Sclater 1893, Gladstone 1927), was a key figure in the history of *Patagona* taxonomy. For many years, Whitely collected specimens in southern Peru, with numerous trips throughout Arequipa and Cusco between the years of 1867 and 1873 (Sclater and Salvin 1867, 1869; Sclater 1893, Chapman 1921, Stephens and Traylor 1983). During this time, Whitely was credited with a number of new species discoveries, including the Bearded Mountaineer (*Oreonympha nobilis*; Sclater 1893). Many of Whitely’s specimens were dispersed through the specimen trading business of his father, Henry Whitely, Sr. (Sharpe 1906, Rasmussen and Prys-Jones 2003). Although Whitely Sr., and to a lesser extent, Whitely Jr.’s, reputations have been questioned (Whitely, Sr. has been called ‘reliably unreliable’; Rasmussen and Prys-Jones 2003), none of the specific problems that have been found appear to concern the localities of Whitely’s *Patagona* specimens.

Whitely collected at least 16 *Patagona* specimens in Peru from 1867 to 1873 (Table S1). In one letter about his 1867 Arequipa trip, Whitely noted, “… now commenced the worst part of the journey…the way, however, was enlivened by the sight of numerous birds, and especially, for some 2,000 feet, by the movements of the Giant Humming-bird (*Patagona gigas*)” (Sclater and Salvin 1867). Whitely’s *Patagona* specimens, particularly those from Tinta, Cusco, Peru, have played a crucial role in history because they are thought to have formed the basis for a subsequent type description by Boucard (Boucard 1893). Whitely collected in Tinta on several trips, including May–June 1868 and January 1869 (Sclater and Salvin 1869). The identities of at least three *Patagona* specimens from his 1868 trip are known (MNHN-ZO-MO-1989-340, FMNH-45797, and FMNH-45798), and three others are suspected (AMNH-37500, AMNH-37501, and MNHN-ZO-MO-1989-34; Table S1). One specimen (AMNH-37501), housed at the AMNH was designated as the “Type of Fig. 33, p. 67” in Giraud Elliot’s 1879 synthesis, “*A classification and synopsis of the Trochilidae*” (Elliot 1879). This specimen, a male from Tinta, was noted in original ledgers as “figured in synopsis”, as was AMNH-37502, a female collected from Cachupata, Cusco, Peru on August 21, 1873; both specimens were denoted with equivalent status (Table 1). These specimens, along with two other Whitely specimens, one each from Cachupata and Tinta (AMNH-37499 and AMNH-37500), were originally part of the Tring Collection of birds but were obtained by the AMNH in 1932 when Lord Rothschild sold his collection (Lanyon 1995; Table S1). Based on available data from gazetteers and related publications, AMNH-37500 and AMNH-37501 are suspected to have been collected during Whitely’s June–August 1868 trip to Tinta; a point of clarification from Williamson et al. (2024). There are at least three other Whitely specimens (ANSP-50447, NHMUK 1887.3.22.700, and NHMUK 1913.3.20.291) collected during January 1869, and one additional specimen (ANSP-50448) is suspected to have been collected in either May–June 1868 or January 1869 (Table S1). At present, Whitely’s *Patagona* specimens are known to reside at the AMNH, MNHN, ANSP, FMNH, and NHMUK; it is possible that there are additional Whitely *Patagona* specimens in other collections, as well.

**Figure S1.**
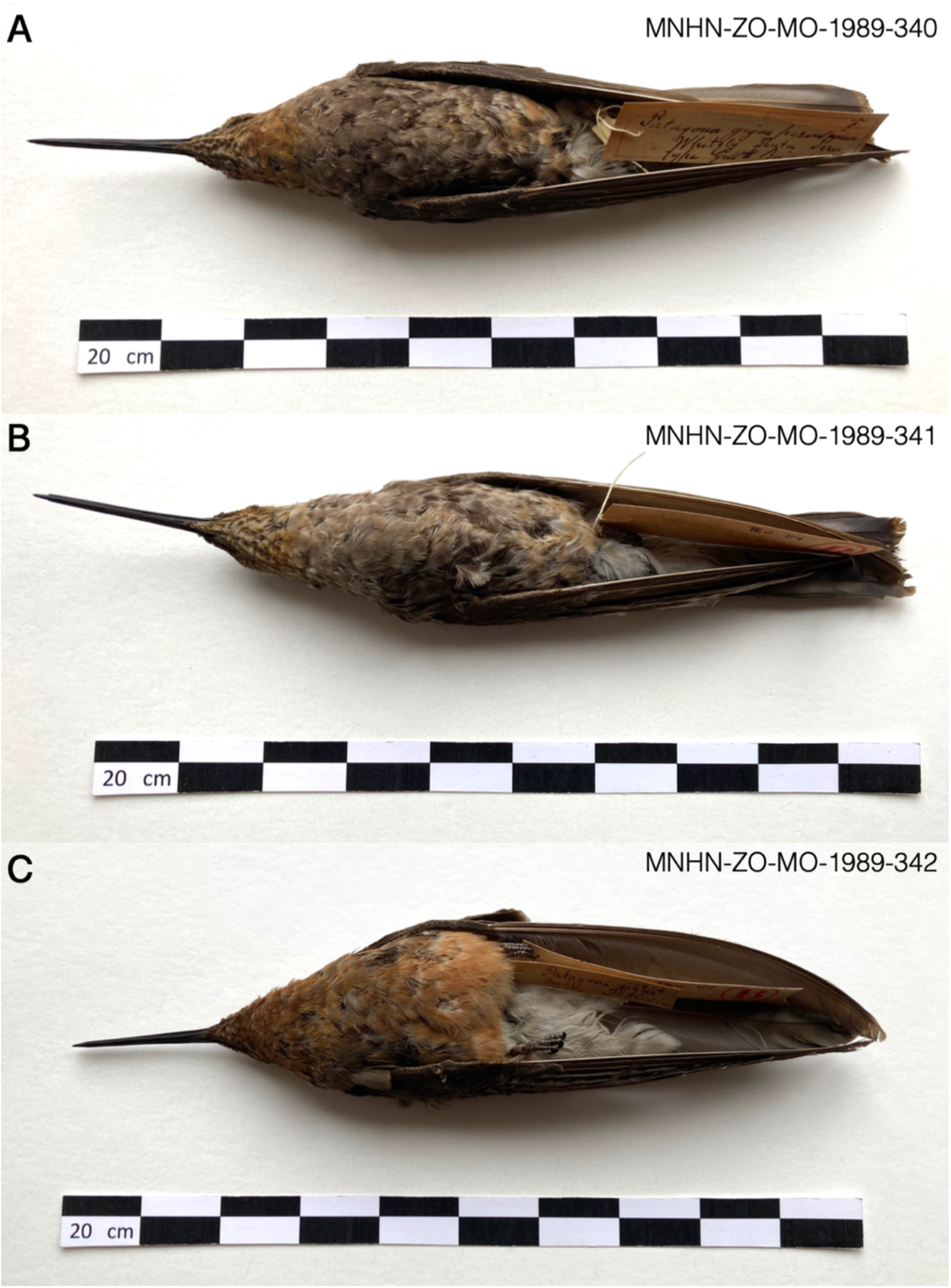
**A** Lectotype for the taxon *peruviana*, MNHN-ZO-MO-1989-340, a male, collected June 15, 1868. **B** Paralectotype for the taxon *peruviana*, MNHN-ZO-MO-1989-341, a female, no date. Both specimens are from Boucard’s (1893) collection and were collected by Henry H. Whitely from Tinta, Peru. **C** Holotype for the taxon *boliviana* from Boucard’s (1893) collection, MNHN-ZO-MO-1989-342, undetermined sex, collected in 1875 by Clarence Buckley from Bolivia. Photos are unedited from originals. Photos: Patrick Bousses, Muséum National d’Histoire Naturelle (MNHN), Paris.

**Figure S2.**
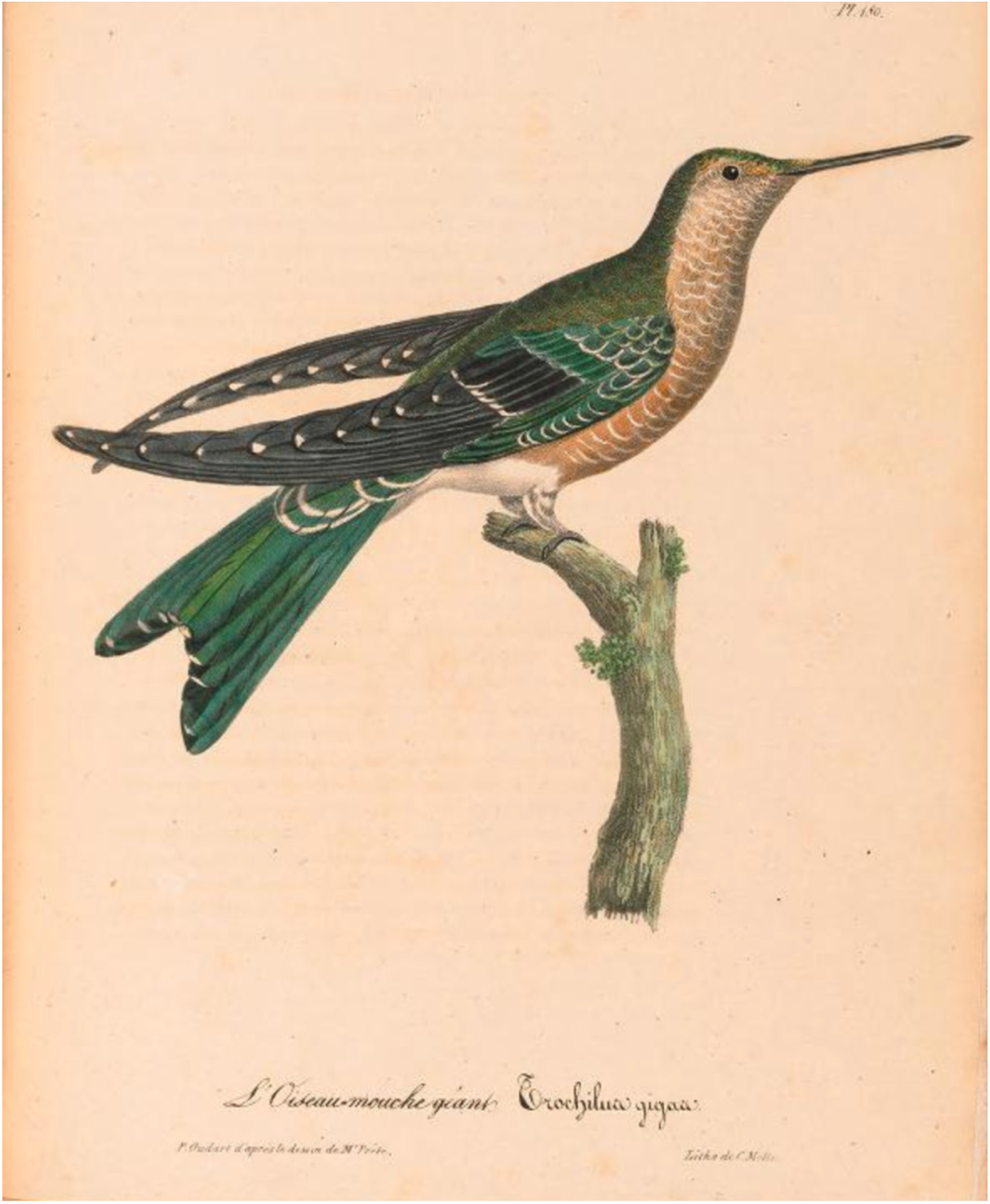
Vieillot’s putative “young male” syntype, illustrated by Paul Louis Oudart in Plate 180 of Vieillot’s *La Galerie des Oiseaux* (Vieillot 1824). The individual depicted was attributed to M. (Mr.; for “monsieur” in French) Portier, “attaché” of the Ministry of the Navy. The specimen has not been located and its current whereabouts are unknown to us.

**Figure S3.**
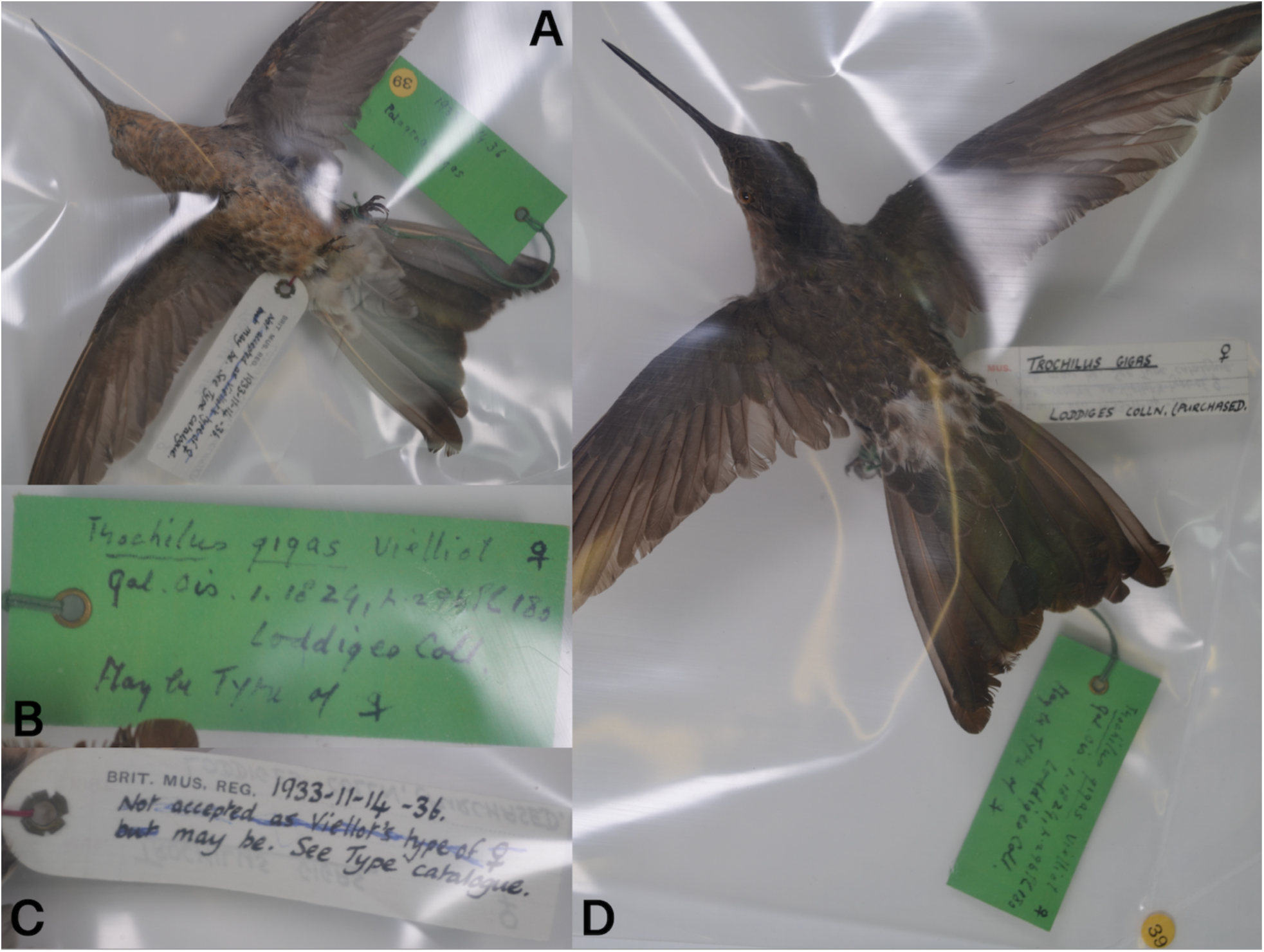
The putative “adult female” syntype of Vieillot (1824) for the species *Patagona gigas*, formerly known as the Giant Hummingbird and currently known as the Southern Giant Hummingbird (*sensu* Williamson et al. 2024). The specimen shown is NHMUK-1933.11.14.36, a female with no collection date and no specific locality (provenance unknown), first acquired by Jean-Baptiste Bécouer before his death in 1777. This specimen, a taxidermied mount, is thought to be one of two syntypes used to describe *Patagona gigas* by Vieillot in 1824; but see specimen tag notes. **A** Ventral view showing the bird’s throat, which is a strong match for the Southern Giant Hummingbird; **B–C** Non-original specimen labels (the specimen does not possess original labels) indicating uncertainty about whether NHMUK-1933.11.14.36 was one of Vieillot’s (1824) syntypes; **D** Dorsal view of the specimen. Photos: Mark Adams, NHMUK.

**Figure S4.**
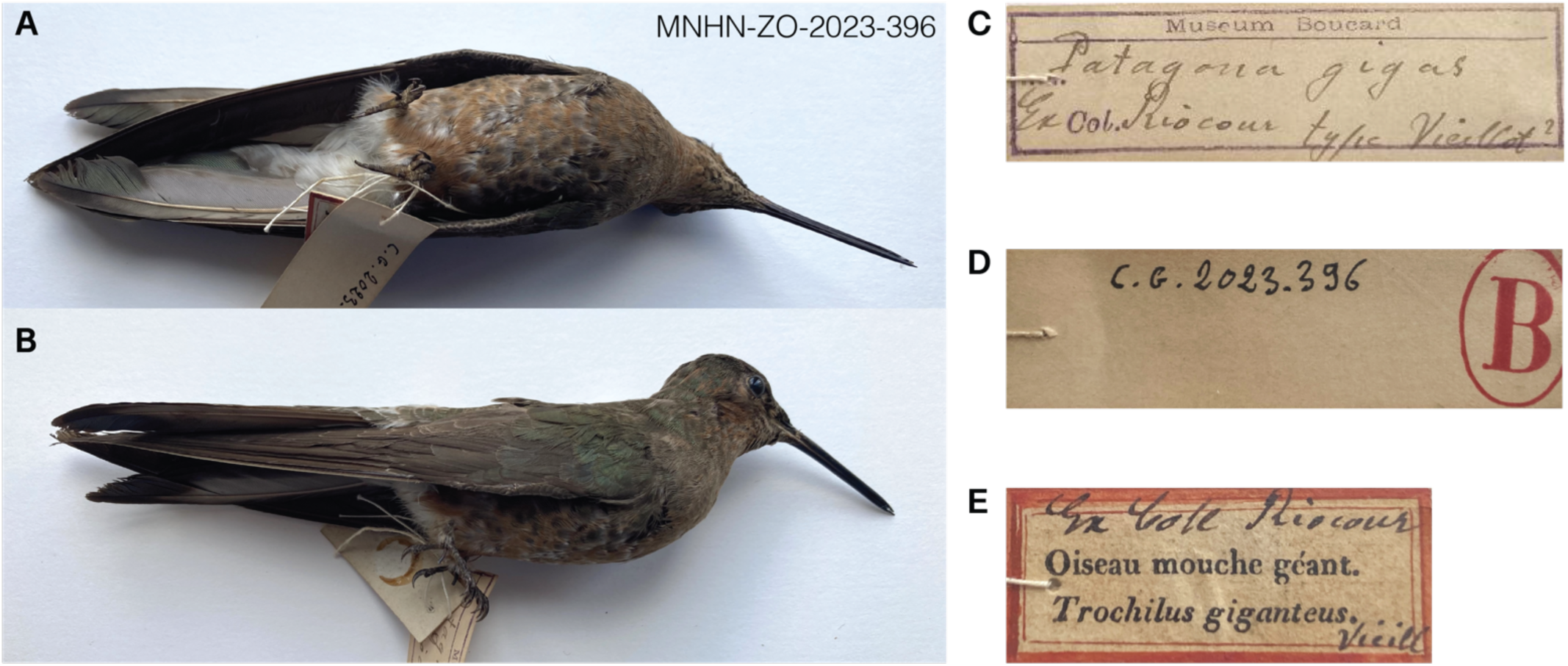
Specimen MNHN-ZO-2023-396 (sex and locality unknown; collection date unknown but prior to 1842) from the MNHN in Paris, believed by Boucard (1893) to be “Vieillot’s type”. Boucard wrote, “I have in my collection what I consider as the type of Vieillot ‘Ex Coll Riocour’” (Boucard 1893). MNHN-ZO-2023-396 is not listed as a syntype by MNHN. **A** Ventral view of specimen. **B** Side view of specimen; photos are unedited from originals. **C** Front side of the Boucard Collection tag showing the question mark (“?”) after the words “type Vieillot” (Patrick Bousses, *pers. comm.*). **D** Back of Boucard Collection tag. **E** Front of Comte de Riocour Collection tag. Photos: MNHN.

**Figure S5.**
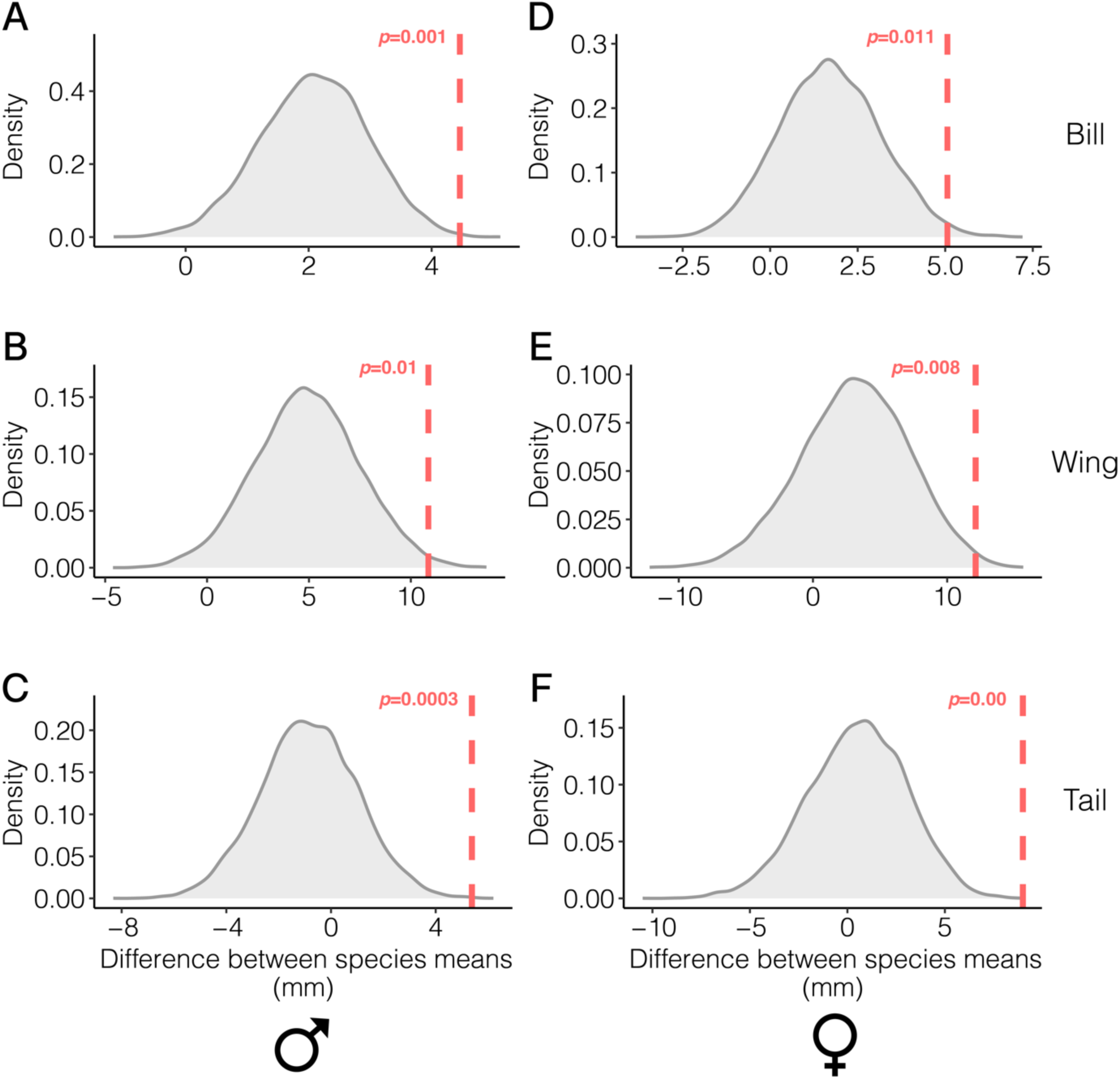
Expected distributions of trait mean differences between Northern and Southern Giant Hummingbirds, based on 10,000 resampled datasets generated from Williamson et al. (present study). **A–C** Expected male trait differences for each bill, wing, and tail; **D–F** Expected female trait differences for each bill, wing, and tail. Red dashed vertical lines indicate the differences between *gigas* and *peruviana* trait means from Hellmayr (1932) data, with the estimated probabilities of obtaining Hellmayr’s values (red text). Estimated probabilities were calculated from 10,000 simulated datasets.

**Figure S5.**
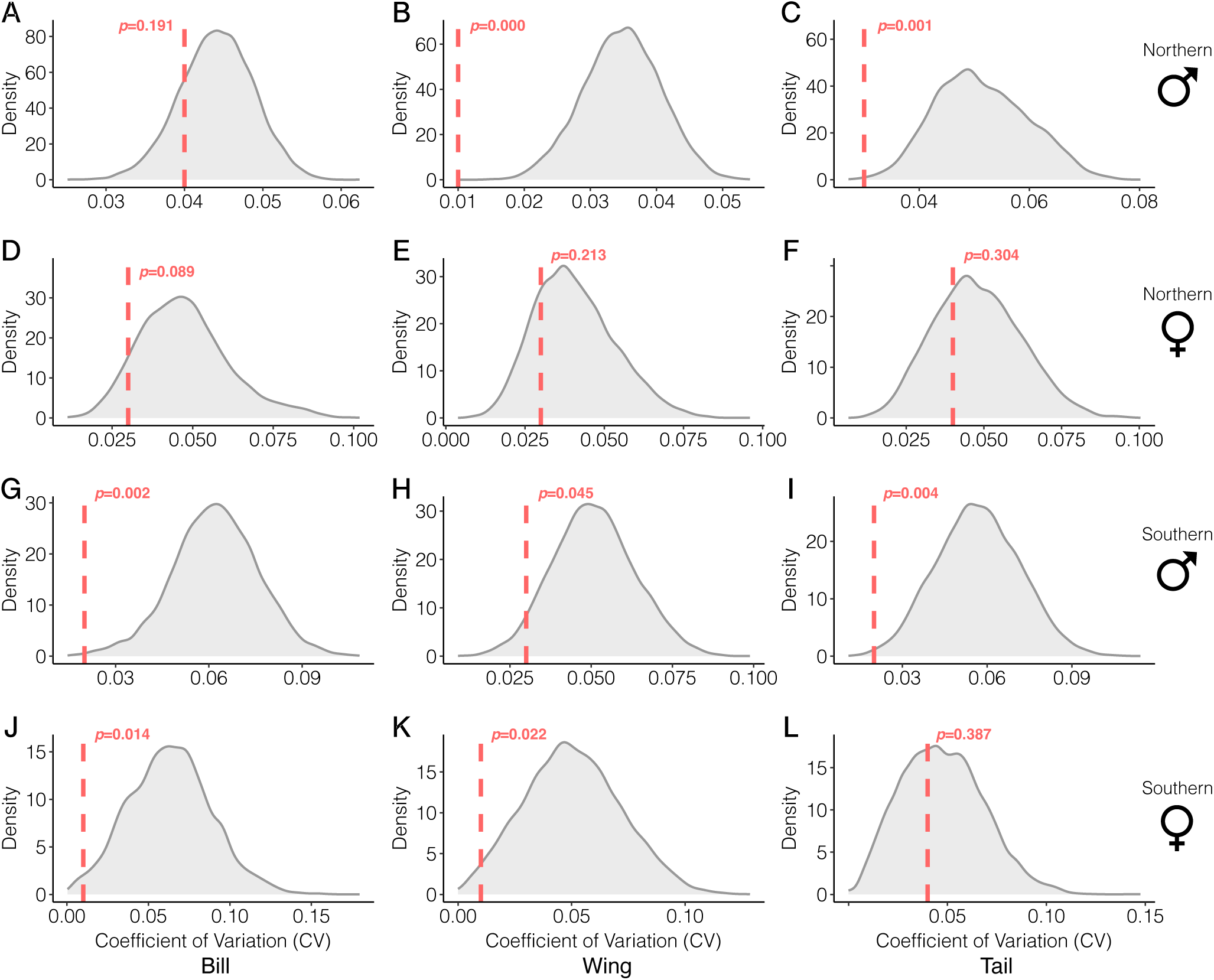
Expected distributions of coefficients of variation (CVs) for each Northern and Southern Giant Hummingbirds, based on 10,000 resampled datasets generated from Williamson et al. (present study). CVs were estimated separately for each species (Northern and Southern Giant Hummingbird), each sex (males and females), and each trait (bill, wing, and tail). **A–C** Northern males; **D–F** Northern females; **G–I** Southern males; **J–L** Southern females. Red dashed vertical lines indicate the CVs from Hellmayr’s (1932) original data, with the estimated probabilities of obtaining Hellmayr’s values (red text). Estimated probabilities were calculated from 10,000 simulated datasets.

**Table S1.**
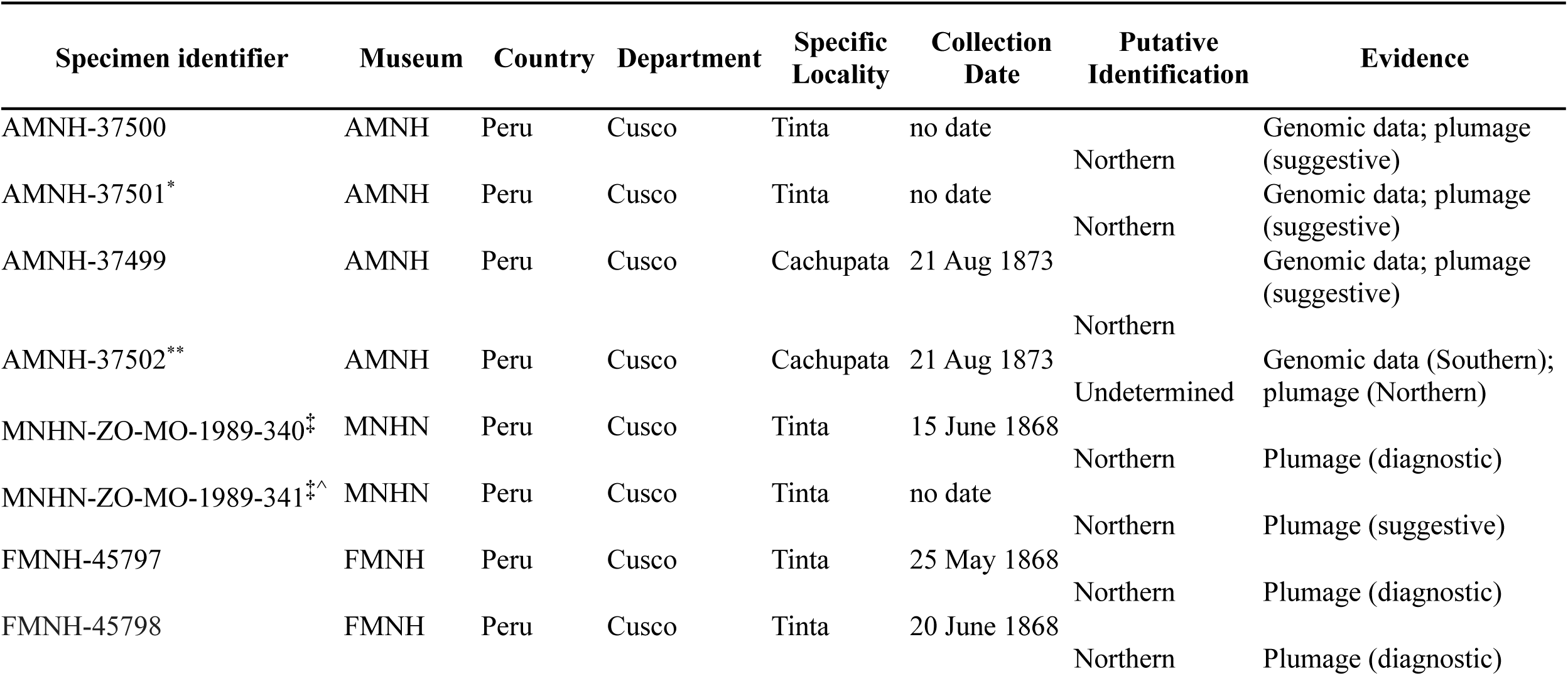
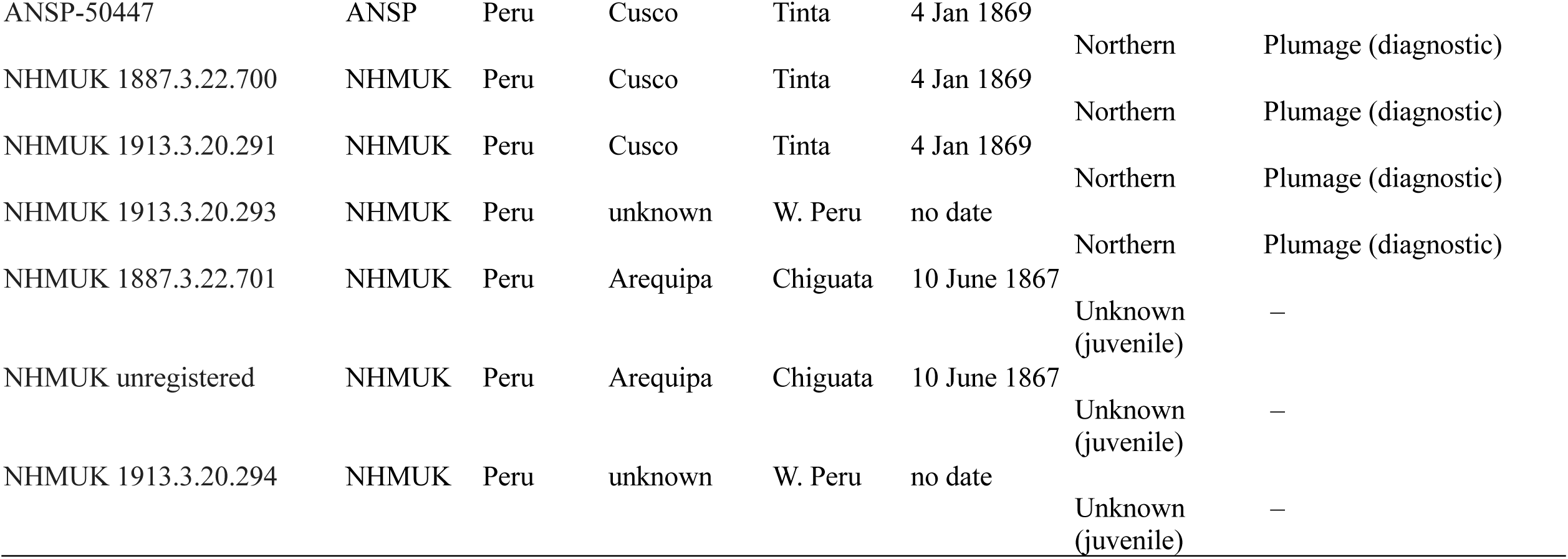
Summary of *Patagona* specimens collected from southern Peru by Henry H. Whitely, Jr. One additional individual (collected ca. July 20, 1867 in Arequipa; known from Whitely’s letter to Philip Lutley Sclater and Osbert Salvin, 1867; (Sclater and Salvin 1867)) is suspected but remains unidentified. Additionally, ANSP-50448 appears to be a Whitely specimen, but its origins and provenance are unknown (Nate Rice, *pers. comm*.) Museums: AMNH, American Museum of Natural History, New York, USA; ANSP, Academy of Natural Sciences of Drexel University, Philadelphia, USA; FMNH, Field Museum of Natural History, Chicago, USA; MNHN, Muséum National d’Histoire Naturelle, Paris, France; NHMUK, Natural History Museum at Tring, Tring, UK. *Indicates status as “type of fig” in Elliot 1879. **Equal status in AMNH ledgers with AMNH-37501. ^‡^Indicates status as a *peruviana* syntype. ^^^MNHN-ZO-MO-1989-341 is listed in the MNHN database as having a collection date of “ca. 1868”, which was estimated based on Whitely’s Peru collection dates. However, the specimen has no date information associated with it, but was likely collected during 1868–1878 (Patrick Bousses, *pers. comm*.), and definitively before Whitely’s death in 1892.

**Table S2.** Morphological summary statistics for *gigas* (Hellmayr 1932) and Southern Giant Hummingbird (Williamson et al., present study) males and females. For each measured parameter, we provide sample size (*n*), mean, standard deviation, range (minimum and maximum values), and coefficient of variation.

| Dataset | Sex | Parameter (units) | *n* | Mean | Standard Deviation | Minimum | Maximum | Coefficient of Variation |
| --- | --- | --- | --- | --- | --- | --- | --- | --- |
| Hellmayr 1932 | Male | Bill | 8 | 34 | 0.76 | 33 | 35 | 0.02 |
|  |  | Wing | 8 | 125.5 | 3.34 | 120 | 129 | 0.03 |
|  |  | Tail | 8 | 80.5 | 1.41 | 79 | 83 | 0.02 |
| Williamson et al. (present study) |  | Bill | 72 | 36.45 | 2.33 | 32.41 | 42.25 | 0.06 |
|  |  | Wing | 69 | 127.81 | 6.71 | 111 | 141 | 0.05 |
|  |  | Tail | 70 | 82.16 | 4.83 | 71 | 92 | 0.06 |
| Hellmayr 1932 | Female | Bill | 4 | 34.25 | 0.5 | 34 | 35 | 0.01 |
|  |  | Wing | 4 | 118.75 | 1.5 | 118 | 121 | 0.01 |
|  |  | Tail | 4 | 75 | 3.37 | 70 | 77 | 0.04 |
| Williamson et al. (present study) |  | Bill | 72 | 36.56 | 2.48 | 30.91 | 41.74 | 0.07 |
|  |  | Wing | 70 | 122.66 | 6.83 | 112 | 139 | 0.06 |
|  |  | Tail | 71 | 79.06 | 4.11 | 71 | 90 | 0.05 |

**Table S3.** Morphological summary statistics for *peruviana* (Hellmayr 1932) and Northern Giant Hummingbird (Williamson et al., present study) males and females. For each measured parameter, we provide sample size (*n*), mean, standard deviation, range (minimum and maximum values), and coefficient of variation.

| Dataset | Sex | Parameter (units) | *n* | Mean | Standard Deviation | Minimum | Maximum | Coefficient of Variation |
| --- | --- | --- | --- | --- | --- | --- | --- | --- |
| Hellmayr 1932 | Male | Bill | 25 | 38.46 | 1.49 | 36 | 42 | 0.04 |
|  |  | Wing | 25 | 136.36 | 2.02 | 133 | 140 | 0.01 |
|  |  | Tail | 25 | 85.88 | 2.33 | 83 | 90 | 0.03 |
| Williamson et al.<br>(present study) |  | Bill | 112 | 38.53 | 1.71 | 34.76 | 42.03 | 0.04 |
|  |  | Wing | 98 | 132.7 | 4.68 | 121 | 147 | 0.04 |
|  |  | Tail | 109 | 81.31 | 4.23 | 66 | 91 | 0.05 |
| Hellmayr 1932 | Female | Bill | 8 | 39.31 | 1.03 | 38 | 41 | 0.03 |
|  |  | Wing | 8 | 130.88 | 3.31 | 126 | 137 | 0.03 |
|  |  | Tail | 8 | 84 | 3.25 | 78 | 87 | 0.04 |
| Williamson et al.<br>(present study) |  | Bill | 108 | 38.26 | 1.89 | 31.93 | 42.76 | 0.05 |
|  |  | Wing | 102 | 125.75 | 5.32 | 109 | 141 | 0.04 |
|  |  | Tail | 108 | 79.57 | 3.98 | 69 | 91 | 0.05 |

**Table S4.** Comparison of differences in taxon-mean trait values between Hellmayr (1932) and Williamson et al. (present study). Hellmayr’s data compare differences between the taxa *peruviana* and *gigas*. Williamson et al. data compare differences between the species Northern Giant Hummingbird and Southern Giant Hummingbird.

| Difference between taxon-mean trait values (mm) |  |  |  |  |
| --- | --- | --- | --- | --- |
| Sex | Source | Bill length | Wing chord | Tail length |
| Male | Hellmayr 1932 | 4.46 | 10.86 | 5.38 |
|  | Williamson et al.<br>(present study) | 2.08 | 4.89 | -0.85 |
| Female | Hellmayr 1932 | 5.06 | 12.12 | 9.0 |
|  | Williamson et al.<br>(present study) | 1.70 | 3.09 | 0.51 |

**Table S5.** List of museum specimens from Peru, Bolivia, Argentina, and Chile that are labeled *P. g. peruviana* (or “*peruviana*” on tags) but that are actually Southern Giant Hummingbirds (*Patagona gigas*). We analyzed the catalog records, tags, and skins of 603 adult and juvenile specimens with locality data from eleven major museum collections. A total of 269 individuals (∼44%) are catalogued with subspecific identifiers; of these, 74 specimens (32.7%) are incorrectly identified as *peruviana* (see Fig. 5). This list is intended to facilitate taxonomic revision within the genus *Patagona* and aid museums in correcting the historical record.

| Specimen identifier | Museum | Country | Department | Specific Locality | Collection Date | Sex |
| --- | --- | --- | --- | --- | --- | --- |
| AMNH-139179 | AMNH | Bolivia | Sucre | Incambo | 20 Nov 1915 | male |
| AMNH-139180 | AMNH | Bolivia | Cochabamba | Totora | 2 Oct 1915 | male |
| AMNH-140718 | AMNH | Argentina | Jujuy | Tilcara | 10 Feb 1916 | female |
| AMNH-140719 | AMNH | Argentina | Jujuy | Tilcara | 9 Feb 1916 | female |
| AMNH-140720 | AMNH | Argentina | Jujuy | Tilcara | 10 Feb 1916 | female |
| AMNH-145020 | AMNH | Peru | Cusco | Huaracondo Canon | 23 July 1916 | female |
| AMNH-148279 | AMNH | Bolivia | Cochabamba | Arque | 5 Feb 1915 | female |
| AMNH-37502 | AMNH | Peru | Cusco | Cachupata | 21 Aug 1873 | female |
| AMNH-479593 | AMNH | Argentina | Tucumán | Lara | 12 Feb 1903 | male |
| AMNH-479594 | AMNH | Argentina | Tucumán | Lara | 12 Feb 1903 | female |
| AMNH-479595 | AMNH | Argentina | Jujuy | Tilcara | 24 Nov 1905 | male |
| ANSP-120834 | ANSP | Bolivia | La Paz | Calacoto | 25 Jan 1935 | male |
| ANSP-120835 | ANSP | Bolivia | La Paz | Calacoto | 24 Jan 1935 | female |
| ANSP-120836 | ANSP | Bolivia | La Paz | Calacoto | 26 Jan 1935 | male |
| ANSP-145414 | ANSP | Bolivia | Cochabamba | Tiraque | 27 Sept 1937 | male |
| ANSP-145415 | ANSP | Bolivia | Potosi | Oploca | 27 Feb 1938 | female |
| ANSP-145416 | ANSP | Bolivia | Potosi | Oploca | 28 Feb 1938 | female |
| ANSP-145418 | ANSP | Bolivia | Potosi | Oploca | 28 Feb 1938 | female |
| ANSP-145419 | ANSP | Bolivia | Potosi | Oploca | 28 Feb 1938 | female |
| ANSP-145420 | ANSP | Bolivia | Potosi | Oploca | 2 Mar 1938 | male |
| ANSP-145421 | ANSP | Bolivia | Potosi | Oploca | 2 Mar 1938 | male |
| ANSP-145422 | ANSP | Bolivia | Potosi | Oploca | 28 Feb 1938 | male |
| ANSP-145423 | ANSP | Bolivia | Cochabamba | Tutimayu | 4 Mar 1937 | female |
| ANSP-145424 | ANSP | Bolivia | Cochabamba | Tutimayu | 3 Mar 1937 | female |
| ANSP-145425 | ANSP | Bolivia | La Paz | Viloco | 1 Apr 1938 | female |
| ANSP-145426 | ANSP | Bolivia | La Paz | Viloco | 31 Mar 1938 | female |
| ANSP-167558 | ANSP | Argentina | Catamarca | Andaluzala | 20 Sept 1917 | male |
| CM-120263 | Carnegie | Bolivia | Cochabamba | Arani | 12 Feb 1927 | female |
| CM-120284 | Carnegie | Bolivia | Cochabamba | Tiraque | 26 Mar 1927 | male |
| CM-52921 | Carnegie | Argentina | Jujuy | Maimara | 17 Nov 1911 | male |
| CM-53156 | Carnegie | Argentina | Jujuy | Maimara | 16 Jan 1911 | male |
| CM-81114 | Carnegie | Bolivia | Cochabamba | Arani | 31 Mar 1920 | male |
| CM-81120 | Carnegie | Bolivia | Cochabamba | Vacas | 2 Apr 1920 | male |
| CM-81121 | Carnegie | Bolivia | Cochabamba | Vacas | 2 Apr 1920 | male |
| CM-81180 | Carnegie | Bolivia | Cochabamba | Cochabamba | 14 Apr 1920 | female |
| CM-81182 | Carnegie | Bolivia | Cochabamba | Cochabamba | 14 Apr 1920 | female |
| CM-81219 | Carnegie | Bolivia | Cochabamba | Mollemolle | 19 Apr 1920 | male |
| CM-81271 | Carnegie | Bolivia | Cochabamba | Mollemolle | 23 Apr 1920 | female |
| CM-81273 | Carnegie | Bolivia | Cochabamba | Mollemolle | 23 Apr 1920 | female |
| CM-81275 | Carnegie | Bolivia | Cochabamba | Mollemolle | 23 Apr 1920 | male |
| CM-81581 | Carnegie | Bolivia | Cochabamba | Cochabamba | 2 Oct 1920 | male |
| CM-81585 | Carnegie | Bolivia | Cochabamba | Cochabamba | 3 Oct 1920 | female |
| CM-94400 | Carnegie | Bolivia | La Paz | Guaqui | 26 Jan 1922 | male |
| CUMV-52570 | CUMV | Argentina | Jujuy | Volcan | 5 Nov 2005 | female |
| FMNH-179394 | FMNH | Bolivia | Cochabamba | Vacas | 15 Mar 1939 | male |
| FMNH-179395 | FMNH | Bolivia | Cochabamba | Vacas | 16 Feb 1939 | male |
| FMNH-179396 | FMNH | Bolivia | Cochabamba | Vacas | 16 Feb 1939 | male |
| FMNH-179397 | FMNH | Bolivia | Cochabamba | Arani | 15 Dec 1940 | male |
| FMNH-179398 | FMNH | Bolivia | Cochabamba | Arani | 15 Dec 1940 | male |
| FMNH-179399 | FMNH | Bolivia | Cochabamba | Arani | 6 Dec 1940 | female |
| FMNH-179400 | FMNH | Bolivia | Cochabamba | Arani | 6 Dec 1940 | male |
| FMNH-179401 | FMNH | Bolivia | Cochabamba | Arani | 6 Dec 1940 | male |
| FMNH-179402 | FMNH | Bolivia | Cochabamba | Colomi | 28 Jan 1938 | female |
| FMNH-179403 | FMNH | Bolivia | Cochabamba | Colomi | 28 Jan 1938 | female |
| FMNH-179404 | FMNH | Bolivia | Cochabamba | Colomi | 28 Jan 1938 | male |
| FMNH-179405 | FMNH | Bolivia | Cochabamba | Colomi | 28 Jan 1938 | female |
| FMNH-179406 | FMNH | Bolivia | Cochabamba | Cochabamba | 15 July 1939 | female |
| FMNH-179407 | FMNH | Bolivia | Cochabamba | Cochabamba | 15 July 1939 | male |
| FMNH-179408 | FMNH | Bolivia | Cochabamba | Tiraque | 5 Aug 1928 | male |
| FMNH-293680 | FMNH | Bolivia | Chuquisaca | Camargo | 4 Feb 1973 | female |
| FMNH-293681 | FMNH | Bolivia | Chuquisaca | Camargo | 5 Feb 1973 | female |
| FMNH-293683 | FMNH | Bolivia | Tarija | Tojo | 25 Jan 1973 | female |
| FMNH-57059 | FMNH | Argentina | Tucumán | Colalao del Valle | 14 Aug 1916 | female |
| FMNH-61681 | FMNH | Chile | Tarapacá | Putre | 3 July 1924 | male |
| FMNH-67517 | FMNH | Peru | Lima | Matucana | 3 May 1922 | male |
| FMNH-67518 | FMNH | Peru | Lima | Matucana | 3 May 1922 | female |
| LSUMZ-35863 | LSUMZ | Bolivia | Cochabamba | Orcoma | 21 July 1960 | female |
| LSUMZ-37404 | LSUMZ | Bolivia | Cochabamba | Cochabamba | 10 Apr 1958 | female |
| MCZ-262196 | MCZ | Argentina | NA | Cumbres de Luracatao | NA Jan 1962 | female |
| MCZ-287952 | MCZ | Chile | NA | Chusmisa | NA Nov 1943 | female |
| MCZ-95880 | MCZ | Bolivia | La Paz | La Paz | NA Sept 1895 | female |
| USNM-172374 | USNM | Bolivia | La Paz | La Paz | 10 Feb 1893 | male |
| USNM-529148 | USNM | Peru | Arequipa | Volcan Misti | 8 July 1938 | male |
| USNM-529149 | USNM | Peru | Arequipa | Volcan Misti | 8 July 1938 | male |

## Notes

### Competing Interest Statement

The authors have declared no competing interest.

### Summary of Updates

Significant updates to main text; additional data and analyses.

